# Dicer1 is reduced in APPswe/PSEN1dE9 mice and is regulated by Nrf2

**DOI:** 10.1101/711572

**Authors:** Yan Wang, Meiling Lian, Jing Zhou, Shengzhou Wu

## Abstract

The pathogenesis of Alzheimer’s disease (AD) involves the central roles of oxidative stress. Oxidative stress due to Dicer1 depletion may underline the neurodegeneration in the central nervous system and degeneration of retinal pigment epithelial cells in geographic atrophy form of age-related macular degeneration. We hypothesized that Dicer1 may play roles in AD pathogenesis. Indeed, Dicer1 was reduced in the hippocampus and cortex of APPswe/PSEN1dE9 mice, an AD model. Dicer1 knockdown induced oxidative stress, mitochondrial dysfunction, apoptosis in cultured neurons, and increased secretions of interleukin-1β/-18, indicators of inflammasome activation. Accordingly, Dicer1 was decreased by amyloid peptide and the effect was connected with down-regulation of nuclear factor erythroid 2-related factor 2 (Nrf2). Anti-oxidant response elements (AREs) were identified in the promoter of *Dicer1* and Keap1-Nrf2-AREs signaling was demonstrated to regulate Dicer1 expression. Furthermore, overexpression of Dicer1 carried by adenovirus in the cultured neurons rescued neurite deficit induced by amyloid peptide. In consistent with the *in vitro* results, injection of Dicer1-overexperssion adenovirus in the hippocampus of the AD mice significantly improved spatial learning. Altogether, we unveiled an unexploited roles of Dicer1 in AD and a novel way of Dicer1 regulation. These findings suggest that Dicer1 may be a target in AD therapy.

**Significance Statement:** Dicer1 is a microRNA-processing enzyme, which is central to microRNA maturation. For the first time, we herein reported that Dicer1 was reduced in the hippocampus or the cortex of AD mice before overt amyloid plque deposition and overexpression of Dicer1 in the hippocampus significantly improved spatial learning in AD mice. We also demonstrated that Dicer1 was regulated by Keap1-Nrf2-ARE signaling which is unreported before. These findings advance understandings of AD pathogenesis and suggest that Dicer1 may be a molecular target in AD therapy.

## Introduction

Alzheimer’s disease (AD) is an inexorable neurodegenerative disorder, characterized with extracellular amyloid plaque deposition, intracellular neurofibrillary tangle formation, and extensive neuronal loss (Hardy and Selkoe, 2002; Mattson, 2004). The pathogenesis of AD involves the key component of oxidative stress which is dependent on the balance between reactive species and anti-oxidation systems (Halliwell, 2006). Accumulation of reactive oxygen/nitrogen species generates oxidative stress and produces ensued cytotoxicity(Markesbery, 1997) as well as does ferroptosis due to iron accumulation in AD brain(Belaidi and Bush, 2016; Stockwell et al., 2017). Amyloid peptide inactivates membrane channel and transporter including sodium/calcium exchanger, calcium ATPase, glutamate and glucose transporter, these hazardous effects involving the roles of ROS and lipid peroxidation(Mark et al., 1995; Keller et al., 1997; Mark et al., 1997). Anti-oxidation systems are used to cope with oxidative injury, in which the regulation of the key enzymes synthesizing glutathione and of the protein supporting the generation and recovery of peroxiredoxins revolves around a transcriptional factor, nuclear factor erythroid 2-related factor 2 (Nrf2)(McWalter et al., 2004; Chen et al., 2005). Nrf2 is a basic leucine zipper protein that regulates expression of a multitude of antioxidant proteins in response to oxidative injuries and is retained in cytoplasm by Keap1 under quiescent situation; upon oxidative stimulation, Nrf2 is released from the binding and translocated into nucleus, triggering the transcription of antioxidant genes(Kobayashi et al., 2004).

Dicer1 is a pre-microRNA processing enzyme which is central to microRNA maturation (Bernstein et al., 2001). Dicer1 is reduced in retinal pigment epithelial cell (RPE) which induces cytotoxicity via Alu RNA accumulation, resulting in oxidative injury via activation of NLRP3 inflammasome (Kaneko et al., 2011; Tarallo et al., 2012); upregulation of NLRP3 inflammasome is also observed in Alzheimer’s brain (Heneka et al., 2013). In addition of the cytotoxicity induced by Dicer1 depletion in the RPEs, Dicer1 loss induces neurotoxicity in different brain regions (Davis et al., 2008; Chmielarz et al., 2017). In light of these evidence, we exploited the roles of Dicer1 in AD and the regulation of Dicer1 expression. We further explored potential therapy by overexpressing Dicer1 in the hippocampus of APPswe/PSEN1dE9 (APP/PS1) mice.

## Materials and Methods

### Materials

The following primary antibodies were used in this study: Dicer1(Sigma, St Louis, MO, USA, cat# WH0023405M1, research resource identifier, RRID: AB_1841286), caspase 3 (Santa Cruz Biotechnology, Santa Cruz, CA, USA, cat# sc-271759, RRID: AB_10709891), Keap 1(Proteintech, Suzhou, China, cat# 60027-1-Ig, RRID: AB_2132623), and rabbit polyclonal anti-Nrf2 (Proteintech, cat# 16396-1-AP, RRID:AB_2782956). Activated caspase 3 (Beyotime Biotechnology, Haimen, China, cat# AC033), βIII-tubulin (Beyotime Biotechnology, cat# AT809), Histone 3 (Beyotime Biotechnology, cat# AF0009), Vimentin (Beyotime Biotechnology, cat# AF1975), GAPDH (Bioworld, Nanjing, China, cat# MB001), NeuN (Beyotime Biotechnology, cat# AF1072). The following secondary antibodies were also used in this study: goat anti-mouse horseradish peroxidase conjugated IgG (Boster Biological Technology, Co. Ltd, Wuhan, China), goat anti-Rabbit horseradish peroxidase conjugated IgG (Boster Biological Technology). The following reagents or cell lines were used in this study: a human Aβ42 peptide (GenScript, Nanjing, China, cat# RP10017), the Dicer1 siRNA duplex and the negative control (NC) siRNA duplex (Genepharma, Suzhou, China), MTS reagent (CellTiter 96 AQueous One Solution, Promega, Beijing, China, cat# 3580), pfu High fidelity enzyme (Qiagen, Beijing, China, cat# KP202), pJet1.2 vector (Thermo Fisher Scientific, Carlsbad, CA,USA, cat# K1231), pGL6-basic (Beyotime Biotechnology, Haimen, China, cat# D2105), pRL-TK Vector (Beyotime Biotechnology, Inc, cat# D2762), a human pCMV-Nrf2 (Sino Biological, Beijing, China, cat# HG17384-U), a mouse pCMV-Nrf2 (Sino Biological, Beijing, China, cat# MG56971-UT), or an empty pCMV3 vector (Sino Biological, Beijing, China, cat# D2602), mitochondrial extraction kit (Beyotime Biotechnology, cat# C3601), 2’,7’-Dichlorodihydrofluorescein diacetate (Thermo fisher, Shanghai, China, cat# 2938), JC-1 probe solution (Sigma, St Louis, MO, USA, cat# CS0760), Neuro-2a (N2A)(ATCC, Manassas, VA cat# CCL-131, RRID:CVCL_0470), SK-N-BE(2) (ATCC Cat# CRL-2271, RRID:CVCL_0528).

### Animal

All mice were fed water and food *ad libitum* in a temperature- and humidity-controlled animal facility with an automatic illumination on a 12-h on/off cycle in Wenzhou Medical University. All experiments and data reports followed the SfN Policy on Ethics and the study was approved by the Animal Care and Use Committee of Wenzhou Medical University (Approval number# wydw2019-0141).

APPswe/PSEN1dE9 mice (Jackson Laboratory, stock number 004462) express a K595N/M586L *Swedish* mutations and a mutant human *presenilin1* with deletion of exon9 under the control of mouse prion promoter. The mice were multiplied and genotyped according to the guidance by Jackson Laboratory. Both genders of 4- and 11-month-old transgenic mice were used and WT littermates were used as controls. Behavioral experiments were performed during the daytime and at the same time of each day.

### Intrahippocampal injection of adenovirus expressing Dicer1

The adenovirus (Ad) incorporating sequences of either Ad-pCMV-EGFP or Ad-pCMV-Dicer1:T2A:EGFP in which the transcription of Dicer1 and EGFP is driven by independent promoter were packaged and generated by Cyagen Biosciences Inc. (Guangzhou, China). Before stereotactic injection, animals were anaesthetized by intraperitoneal injection of ketamine/xylazine (0.1/0.05 mg/g body weight) and mounted on a stereotactic frame (KOPF, KD Scientific). The virus were injected into bilateral hippocampus with 2µl of viral titre (1.2 × 10^9^ vg/mL) in each hemisphere using a 10-µl Hamilton syringe (Hamilton Medical, Reno, NV, USA) connected to a 30-gauge micropipette at an injection rate of 0.2 µl/min. The injection coordinates were anteriorposterior, −2 mm, mediolateral, ± 2mm, dorsoventral, −2mm from bregma. The person in charge of injection wore protective equipment and the injections were conducted in a biological safety cabinet in a biological safety level II lab setting. Autoclave of disposals was used to prevent contamination. The injections were conducted in 3.5-/4-month-old mice and behavior test was performed during 17-23 days after injection in which learning curve was acquired during 17-22 days and probe trial conducted at the 23rd day after injection. Halves of mice were used for immunohistochemistry and halves of them used for western blot.

### Morris Water Maze

Water maze tests were performed using three groups of mice, WT mice (with intrahippocampal injection of Ad-EGFP) and APP/PS1 mice (intrahippocampal injection of Ad-EGFP or Ad-Dicer1-T2A:EGFP). The water maze (1.2-m diameter) was filled with water (24°C) and made opaque by the addition of nontoxic white paint. The water maze was surrounded by a black curtain (placed 80 cm away) that held three salient visual cues. Initially, mice were randomly trained in four quadrants of the water maze, and were allowed a maximum of 60 s to find a hidden platform (10-cm diameter, 1 cm under the water surface). Mice were trained for 6 days (4 trials per day, 10-min interval between trial). On day 7, mice were given a 60-s probe test (scanning the platform) to test their spatial memory. SLY-WMS Morris Water Maze System (Beijing Sunny Instruments Co. Ltd) consisting of a frame grabber, a video camera, a water pool, and the analysis software was used to monitor the animal’s swimming pattern, distance, speed, and the amount of time spent in each of the four quadrants.

### Cell culture

Newly born mice were used for culturing dissociated cortical and hippocampal neurons (CNs and HNs, respectively). Cortex or hippocampal tissue was mechanically dissociated after digestion with 0.25% trypsin (Beyotime, Haimen, China) and 0.1% DNaseI (Takara, Dalian, China) at 37°C for 15 min. DMEM/F12 containing 10% fetal bovine serum (Gibco, Billings, MA,USA) was then added, and the cells were centrifuged at 120 g for 3 min and the supernatants were then plated on poly-D-lysine-coated 8 cm-culture dishes in neurobasal medium supplemented with 2% B27 (Invitrogen, Carlsbad, CA,USA) and 10 μM cytosine arabinoside (Sigma-Aldrich, Saint Louis, MO, USA). After 24 hours, the culture medium was replaced with fresh neurobasal medium supplemented with 2% B27 and the neuronal cultures were used for experiments after three days *in vitro culture* (DIV 3). Before transfection, the medium was replaced with DMEM/F12 containing 0.1% bovine serum albumin. After 24 h in culture, neurons were used for transfection, Aβ42 oligomer treatment or adenovirus infection. N2A, SK-N-BE(2) or HEK 293T cells was cultured in Dulbecco’s modified Eagle’s Medium(DMEM) containing 10% Fetal bovine serum(FBS).

### Plasmid construction and Luciferase reporter assay

With genomic DNA extracted from HEK 293T cells as a template, the region upstream of Dicer1 transcription start site (tss) (Entrez acc.no.:NM_177438.2) were amplified, producing an amplimer containing sequence from −1230 (relative to tss) to +3 (promoter 3), using forward primer CGACGCGTTTGC**AGTGAGC**TGAGATTGCGCCAC-TACATT and reverse primer CCCAAGCTTCCCAAGCTTCCCAAGCTTCTTTGTGTCC. Two more regions were amplified spanning −1131 or −756 to +3 by alternating forward primer CGACGCGTAGCCTGGG**TGACAGAGC**GAAACTCT (promoter 2) or CGACGCGTACC**TGAGCTGGC**TGGGACCCAGCATTTA (promoter 1), respectively. The bold letters indicated the predicted AREs sites with consensus sequence TGA(C/G)NNNGC. A sequence not containing AREs site was also amplified by changing a forward primer CGACGCGTGGGCAGTCAGAGAGAGAGGAAAGGAAGG spanning −581 to +3 (promoter 0). The region spanning individual ARE sequence was also amplified by using forward primer CGACGCGTAGGAGATCAAGACCATCCTGGG and reverse primer CCCAAGCTTTAGTGGCGCAATCTCAGCT for ARE3 (124 bp). ARE2 (192 bp) was amplified using forward primer CGACGCGTCAGCCTGGGTGA CAGAGCGAAAC and reverse primer CCCAAGCTTCTGGTCTGCAAGGCA. ARE1(185 bp) was amplified using forward primer CGGCACGCGTGAGCTGCC TTGTGACTTTGCCTTC and reverse primer CCCAAGCTTCGTCTTTCAACACTTGA -TCA. PCR products were amplified by pfu High fidelity enzyme and cloned into a pJet1.2 vector and then subcloned into pGL6-basic containing firefly luciferase coding sequence. The constructs spanning tss were referred to as Dicer 1 promoter 3-Luc, promoter 2-Luc, promoter 1-Luc, and promoter 0-Luc and the constructs containing individual AREs were referred to as ARE3-Luc, ARE2-Luc, and ARE1-Luc. ARPE1 in promoter 1-Luc was mutated with forward primer incorporating mutated base pair in bold letters **GT**ACCTGGCTGGGACCCAGCATTTA and reverse primer CCCAAGC-TTCCCAAGCTTCTTTGTGTCC; promoter 1-Luc was used as the template. Using the resultant plasmid as the template, the forward primer incorporating Mlu I restrict site and the reverse primer containing HindIII restriction site were used to generate product, which was subcloned into PGL6-basic, namely, mutant promoter-1. The forward primer was CGACG<u>CGTACC</u>**GT**ACCTGGCTGGGACCCAGCATTTA and the reverse primer was CCC<u>AAGCTT</u>CCCAAGCTTCCCAAGCTTCTTTGTGTCC with restriction sites underlined.

The HEK293T cells were seeded at a density of 3 × 10^4^ cells/per 48 wells and transfected according to the manufacturer’s protocol (Lipofectamine 2000, Invitrogen). The *Renilla* luciferase-containing reporter (pRL-TK Vector) was mixed with individual construct containing firefly luciferase, and a human pCMV-Nrf2 or an empty pCMV3 vector at a ratio of 1:20:20, respectively. The relative luminescence unit (RLU) of luciferase/renilla assay was measured per the manufacturer’s protocol (Beyotime Biotechnology, cat#RG027) by a plate reader (SpectraMax M5, Molecular devices, San Jose, CA, USA).

### Aβ42 oligomer preparation

The oligomer form of Aβ42 was prepared as described (Rushworth et al., 2013). Briefly, the peptide was dissolved in 1,1,1,3,3,3-Hexafluoro-2-propanol (Sigma-Aldrich) to remove any aggregates, stored in 4°C for 30 min, dried under room temperature, and dissolved in dimethyl sulfoxide (DMSO, Sigma-Aldrich) to 1 mM. The dissolved peptide was diluted in phenol-free F12 medium to 500 nM and incubated at 4°C for 24 h. The solution was subjected to centrifugation at 14,000 x g for 30 min at 4°C. The supernatant was collected as the oligomer preparation which was used at a final concentration of 100 nM.

### Cell viability assay

Neurons were cultured in 96-well plates at a density of 5 X 10^3^ per well in DMEM/F-12 medium supplemented with B-27. After transfected with Dicer1 siRNA duplex (sense, 5’-GCACAUCAAGGUGCUACUATT-3’, antisense, 5’UAGUAGCACCUUGAUGUGCTT-3’) or negative control siRNA duplex (sense, 5’-UUCUCCGAACGUGUCACGUTT-3’, antisense, 5’-ACGUGACACGUUCGGAGAATT-3’) by lipofectamine 2000 for 12 h, the medium was replaced with fresh DMEM/F-12 medium and continued to culture for 36 h. At the end of treatment, each well was added with 10 µL MTS reagent (500 µg/mL) for 4 h. The absorption values were read in a plate reader at 490nm (SpectraMax M5, Molecular devices).

### Measurement of mitochondrial membrane potential and reactive oxygen species (ROS) activity

Primary cultured cortical neurons at 9×10^5^ cells/well and hippocampal neurons at 5×10^5^ cells/ well were plated into 6-well plates and cultured until DIV3. The neurons were transfected with Dicer1 duplex siRNAs (50 pM) per well by lipofectamine 2000 for 12 h in DMEM/F12 plus 2% B27. After transfection, the cultured medium was replaced with fresh neurobasal medium supplemented with 2% B27 and continued to culture for 36 h. For mitochondrial membrane potential(ΔΨm) assay, the neurons at the end of treatment were stained with JC-1probe solution at 37 °C for 20 min and observed under fluorescence microscope (DMi8, Leica Biosystems, Wetzlar, Germany). The mitochondria were also extracted by a mitochondrial extraction kit. Neural mitochondrion (30 µg) from each group was incubated with JC-1 probe solution at 37 °C for 20 min. The reaction product was added into a 96-well plate for fluorescence measurement, which was read with excitation at 490 nm and emission at 520 nm for monomer (green) and with excitation at 530 nm and emission at 590 nm for aggregates (red). The ΔΨm values were represented by the ratios between aggregate and monomer fluorescence in relative fluorescence units (RFU). For reactive oxygen species (ROS) assay, CNs and HNs were incubated with 10 µM 2’,7’-Dichlorodihydrofluorescein diacetate in neurobasal medium supplemented with 2% B27. After incubation at 37°C for 30 min, the neurons were added into 96-well plate at the density of 4×10^4^ per well and subjected to measurement with plate fluorescence reader at 485nm (SpectraMax M5, Molecular devices).

### Enzyme-linked Immunosorbent assay for secreted IL-1β and IL-18

The primary cultured neurons were seeded into 6-well plate at 5×10^5^ per well. The neuronal cultures were treated with oligomer Aβ42 at 100 nM for 24 h, and then infected with Ad-Dicer1-T2A:EGFP or vehicle virus Ad-EGFP at 2×10^8^ vg/mL per well for 48 h. The culture medium were collected and examined by IL-1β or IL-18 enzyme-linked immunosorbent assay kit following manufacturer’s instruction (BOSTER Biological Technology Co. Ltd, Wuhan, China).

### Neurite outgrowth assay

The neuronal cultures were treated by oligomer Aβ42 at 100 nM for 24 h and then infected with Ad-EGFP or Ad-Dicer1-T2A:EGFP virus (5 X10^7^ vg/mL) for 48 h. For determination of neurite, the neurites at least two times longer than the soma diameter were included in the measurement (Image pro plus, Olympus Optical Co., Ltd, Tokyo, Japan). The measurement of neurite length was conducted in three independent culture preparation and the values were averaged from 150 infected neurons in either group.

### Western blot analysis

The 4- or 11-month-old mice were fasted overnight, and at the next morning, mice were deeply anesthetized by isoflurane and transcardially perfused with ice-cold phosphate buffer (PBS). The hippocampi were isolated and freshly frozen in liquid nitrogen. For analyzing the neurons treated by amyloid peptide, the cytoplasmic and nuclear proteins were extracted by plasma and nuclear protein isolation kit (Cat# P0027, Beyotime Biotechnology), respectively. For analyzing the cells for overexpressing Nrf2 plasmid, SK-N-BE (2) or N2A cells were seeded in 6-well plates at the density of 1×10^6^ or 5×10^5^ cells/well, respectively. Tissue or cell samples were homogenized in Super RIPA buffer composed of components in mM: 460 Tris–HCl, pH 7.4, 138 NaCl, 1 EDTA, 2.5 NaF, 2.5 Na3VO4, 1 phenylmethanesulfonylfluoride, 1 dithiothreitol, supplemented with 0.1% Nonidet P-40, and 1 X protease/ 1 X phosphatase inhibitor cocktail (Sigma, St.Louis, MO). Proteins were resolved on 12% SDS-PAGE gels (Bio-Rad) and transferred to nitrocellulose membranes (ThermoFisher). Membranes were blocked in blocking buffer (Beyotime Biotechnology) and then incubated with primary antibody. Following wash, the membranes were then incubated with secondary antibody, and developed with chemiluminescence system kit (ThermoFisher). The blots were quantified using Image J software (National Institutes of Health, Bethesda, MD, USA).

### Immunofluorescence

The 4- or 11-month-old WT, APP/PS1 non-fasted mice were deeply anesthetized by isoflurane, killed, and then transcardially perfused with ice-cold PBS followed by 4% paraformaldehyde (PFA) in PBS. The isolated brains were postfixed in 4% PFA overnight, then stored in PBS containing 0.01% sodium azide. Brains were mounted in agarose and 10 µm coronal sections were obtained using a Leica VT1000S Vibratome. Antigen demasking was performed in 0.01M sodium citrate solution (pH 6.0) at 99°C for 40 min. After washing the sections in PBS three times, 5 min each, at room temperature (RT), the sections were blocked using 5% normal donkey serum in PBS containing 0.2% Triton X-100 for 1 h at RT. The sections were incubated with the rabbit anti-NeuN (1:800, Beyotime Biotechnology) or monoclonal anti-Dicer 1(1:300, Sigma-Aldrich) overnight at 4°C. For mice injected with virus, the sections were incubated with anti-Dicer 1(1:300, Sigma-Aldrich) or anti-activated caspase 3 (1:300, Beyotime Biotechnology). After washing 3 times with PBS (3×5 min at RT), sections were incubated with secondary antibodies: Alexa Fluor 594 goat anti-rabbit or Alexa Fluor 488 goat anti-mouse IgG (1:1000, Life Technologies Corporation, Carlsbad, CA, USA) for 1 h at RT, then washed three times with PBS (3×5 min at RT). The sections were then mounted on coverslips (Citotest Labware Manufacturing Co., Ltd, Jiangsu, China) and the images of CA3 hippocampus were obtained with a fluorescent microscope (DMi8, Leica Biosystems).

### Experimental design and statistical analysis

During Water Maze test, mice were evenly allocated to each experimental group according to the gender and ages and the tests were conducted in a blinded manner, that is, the investigator was blinded to the group allocation of the genotype or treatment during the experiment and when assessing all the results. Animal experiment sample size calculator: http://www.lasec.cuhk.edu.hk/sample-size-calculation.html was used to determine the sample sizes, assuming 40% difference in the mean and standard deviation of 20% at power 95% and type I error 0.01 (Jiang et al., 2017). Similarly, determination of neurite outgrowth was conducted in a blinded manner. All the data acquired during experiments were included for analysis and there were no sample size differences between the beginning and end of experiments. All data are presented as means ± SEM unless specified. Single data points were shown as overlaying dot-plot on bar-graph when sample sizes were smaller than 15, otherwise, shown as bar-graph. For parametric data, Student’s *t-test* was used for comparing differences between two groups. In experiments with more than two groups, one-way ANOVA was performed followed by Tukey’s *post hoc* test for comparisons among groups. For analysis of two groups of non-parametric data, the Mann-Whitney *U*-test was used for determining the differences. For Morris water maze experiment, two-way ANOVA with repeat measures followed by Tukey’s *post hoc* test was used for analyzing the time spent in four quadrants, frequency to crossing platform, swimming speed and distance. Statistical analyses were performed with Graphpad Prism 7.04 software (Graphpad software, Inc. La Jolla, CA, USA). Difference was regarded as significance when *p*<0.05.

## Results

### Dicer1 was reduced in the hippocampus and cortex of APP/PS1 mice and Dicer1 knockdown induced neurotoxicity

Dicer1 is reduced in the retinal pigment epithelial cells in geographic atrophy form of age-related macular degeneration(Kaneko et al., 2011) and Dicer1 knockout induces neurodegeneration (Davis et al., 2008; Shin et al., 2009). In view of these evidence, we explored Dicer1 expression in AD mice, APP/PS1. Dicer1 staining was significantly reduced in the CA3 hippocampus of APP/PS1 mice compared to WT littermate at the age of 4 months or 11 months (Figure.1A). The protein was also significantly reduced in the homogenates from the hippocampus of 4-month APP/PS1 mice compared to WT littermate (p=0.0092, n=6 for each genotype, Mann–Whitney *U*-test) and this reduction became severe at the age of 11 months (p=0.0002, n=6 for each genotype, Mann–Whitney *U*-test) (Figures.1B and 1C). Similarly, Dicer1 was significantly reduced in the homogenates from the parietal lobe of APP/PS1 mice at the age of 6 months when compared to WT littermate (p=0.022, n=6 for each genotype, Mann–Whitney *U*-test)(Supplemental Figure.1). Nrf2 is a master regulator of anti-oxidation genes regulating redox homeostasis(Rada et al., 2012); Dicer1 depletion induces cytotoxicity via oxidative stress(Kaneko et al., 2011). Thus, we simultaneously detected the protein levels of Nrf2 and activated caspase3 in the homogenates from APP/PS1 hippocampi. Similarly, Nrf2 protein levels in AD mice were significantly reduced compared to WT littermate at the age of 4 months (p=0.0001, Mann–Whitney *U*-test) and 11 months (p=0.0038, Mann–Whitney *U*-test), respectively (Figures. 1B and 1D). Activated caspase3 in AD mice was significantly increased compared to WT littermate at the age of 4 months (p=0.0098, Mann–Whitney *U*-test) and 11 months (p=0.0008, Mann–Whitney *U*-test), respectively (Figures. 1B and 1E).

**Figure 1.**
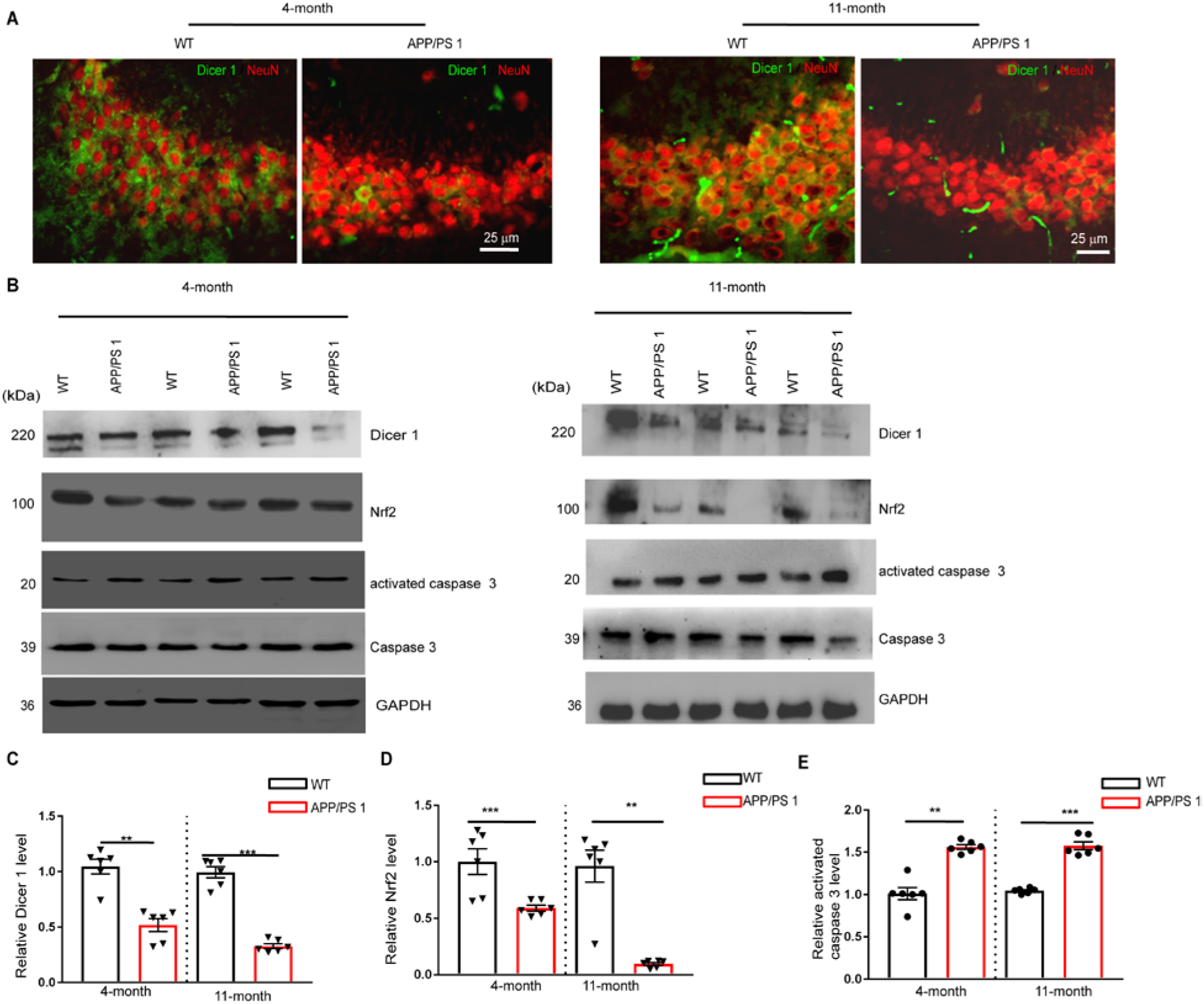
Reduction of Dicer 1, decreased Nrf2 and increased activated caspase-3 in the hippocampus of APP/PS1 mice. (A) APP/PS1 brains were processed into sections which were doubly stained for Dicer1(1:300) (green) or NeuN (1:800) (red). The images were typical staining of CA3 hippocampus (n=3 mice per genotype). Bar, 25 μm for all panels. **(B)** The hippocampi at the age of 4-/11 months were isolated from the brain, homogenized, and subjected to centrifugation at 12,000 X g for 10 min. The supernatants were collected and subject to examination of Dicer1(1:600), Nrf2 (1:600), caspase-3 (1:500), and activated caspase-3 (1:600) by western blot. GAPDH (1:1500) was used as a loading control. The protein levels of Dicer1 and Nrf2 in the hippocampal tissues were reduced in APP/PS1 mice (n=6) compared to WT littermate (n=6), respectively, at the age of 4 months (Left panel) or at the age of 11 months (right panel). By contrast, the protein levels of activated caspase-3 were increased in APP/PS1 mice (n=6) compared to WT (n=6) at the age of 4-/11 months. **(C)** The optical densities of Dicer1 relative to GAPDH were normalized to WT at the age of 4-/11 months (Mann-Whitney U test, **p=0.0092 in 4-month; ***p=0.0002 in 11-month, n=6 per genotype). **(D)**, Nrf2 relative to GAPDH were normalized to WT (Mann-Whitney U test, ***p=0.0001 in 4-month, **p=0.0038 in 11-month, n=6 per genotype). **(E)** Activated caspase 3 relative to GAPDH were normalized to WT (Mann-Whitney U test, **p=0.0098 in 4-month, ***p=0.0008 in 11-month, n=6 per genotype), respectively.

Dicer1 depletion induced cytotoxicity involving the roles of ROS and inflammasome in the RPE cells (Tarallo et al., 2012). Thus, we explored the relevant effects by knocking down Dicer1(Dicer1 siRNA), the effect compared to the cells transfected with negative control scrambled siRNA (NC siRNA) in the primary murine cultures of cortical neurons (CNs, t(2)=19.86, p=0.0025, paired *t*-test) or hippocampal neurons (HNs, t(2)=4.606, p=0.044, paired *t*-test) (Figure. 2A). As expected, Dicer1 knockdown by its specific siRNA induced ROS production in CNs (t(7)=9.539, p=0.00001, paired *t*-test) or in HNs (t(7)=21.77, p=0.00001, paired *t*-test) compared to NC siRNA (Figure. 2B). Similarly, Dicer1 knockdown reduced mitochondrial membrane potential indicating by JC-1 staining in CNs (t(2)=5.085, p=0.0369, paired *t*-test) or in HNs (t(2)=4.853, p=0.0399, paired *t*-test) (Figures. 2C and 2D). Under expectation, Dicer1 knockdown increased the contents of interleukin-1β(IL-1β) in CNs (t(2)=4.978, p=0.0381, paired *t*-test) or in HNs (t(2)=4.907, p=0.0391, paired *t*-test) (Figure. 2E) and the production of IL-18 in CNs (t(2)=5.361, p=0.0331, paired *t*-test)or in HNs (t(2)=11.86, p=0.0070, paired *t*-test) (Figure. 2F), indicating inflammasome activation. We further explored the effect of Dicer1 knockdown on neuronal survival and its related signaling. Under the condition of Dicer1 knockdown, neuronal survival was compromised in CNs (t(2)=4.863, p=0.0028, paired *t*-test) or in HNs (t(2)=12.1, p=0.0012, paired *t*-test) (Figure. 2G) and the protein levels of activated caspase3 were also increased in the CNs (t(2)=11.09, p=0.00080, paired *t*-test) or HNs (t(2)=6.965, p=0.0020, paired *t*-test) comparing to the effects of NC siRNA (Figures. 2H and 2I).

**Figure 2.**
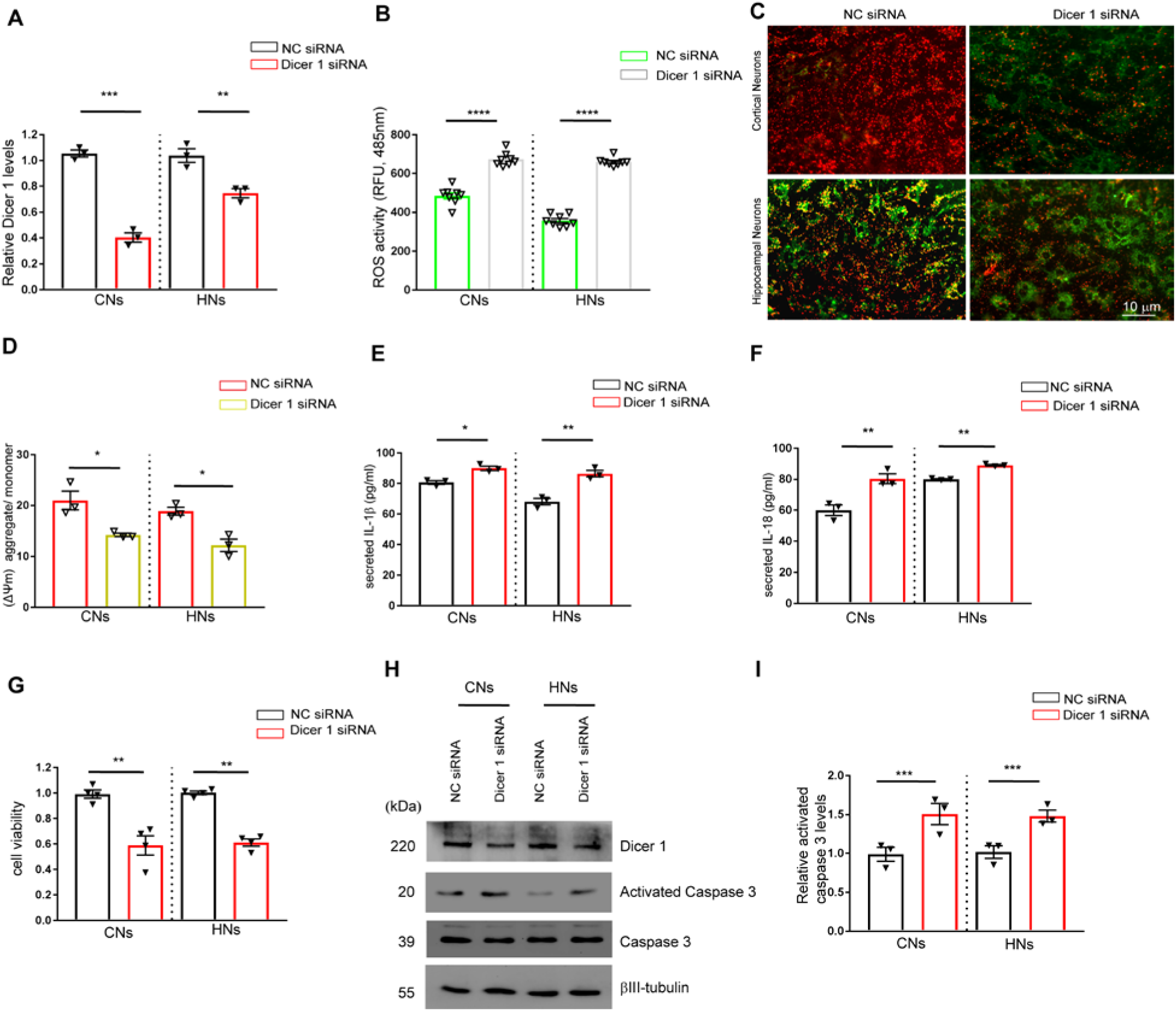
knocking down Dicer 1 induced production of reactive oxygen species (ROS), reduced mitochondrial membrane potential, enhanced secretion of IL-1β, IL-18, and apoptosis in primary cultured neurons. **(A)** Primary murine cortical neurons (CNs) and hippocampal neurons (HNs) in 6-well plates were transfected with Dicer1 siRNA (50 pM) or control siRNA duplex (50 pM) per well by lipofectamine 2000 for 12 h in DMEM/F12 plus 2% B27. The medium was replaced with fresh neurobasal supplemented with 2% B27 and continued to be cultured for 36 h. The effects of knockdown by Dicer1 siRNA were normalized to those of NC siRNA and averaged from three independent culture preparation. Student’s *t*-*test* was used to compare the differences, **p=0.0025 for comparison in CNs and *p=0.044 for comparison in HNs for Dicer 1. (**B**) The ROS levels in CNs or HNs were measured by incubating 2’,7’-dichlorodihydrofluorescein diacetate for 20 min at 37 °C. The fluorescence values in relative fluorescence units (RFU) were acquired in a plate reader at 485 nm and were averaged from four independent culture preparation with duplicate cultures in each preparation. Student’s *t-test* was used to compare the differences of ROS production in neurons between transfection with Dicer1 siRNA and control siRNA. ****p=0.00001 in CNs, ****p=0.00001 in HNs. **(C)** Representative images of JC-1 staining in CNs and HNs subject to Dicer1 knockdown as above, Bar 10 µm for all panels. **(D)** Mitochondrial membrane potential (Δm) of CNs and HNs subject to Dicer1 knockdown. The neurons were subject to Dicer1 knockdown thereof and mitochondria (30 µg) were isolated and stained with JC-1. The values of mitochondrial membrane potential (Δm) were indicated by the ratios between aggregated/monomer RFU and averaged from three independent culture preparations. Student’s *t-test* was used to compare the differences, *p=0.0369 in CNs, *p=0.0399 in HNs. **(E, F)** secreted IL-1β and IL-18 in CNs and HNs subject to Dicer1 knockdown. The CNs or HNs were subjected to Dicer1 knockdown as above. The culture media (100 µL) from neuronal culture subjected to transfection with Dicer1 siRNA or control siRNA duplex were collected and measured by ELISA. The concentrations of IL-1β and IL-18 were averaged from three independent culture preparation. Student’s *t-test* was used to compare the differences. *p=0.0381 when comparing IL-1β production in CNs and *p=0.0391 in HNs. *p=0.0331 when comparing IL-18 production in CNs and **p=0.007 in HNs. **(G)** Viability of neurons subjected to Dicer1 knockdown. CNs or HNs at the density of 5×10^3^ were transfected with Dicer 1 siRNA and NC siRNA for 12 h in DMEM/F12 plus 2% B27, which was replaced with fresh neurobasal medium supplemented with 2% B27 and continued to be cultured for 36 h, respectively. The absorption values at OD 450nm in Dicer1 siRNA group were measured and normalized to NC siRNA group. Student’s *t-test*, **p=0.0028 for comparison in CNs, **p=0.0012 for comparison in HNs. **(H)** Representative images of Dicer 1, caspase3 and activated caspase3 in CNs and HNs subject to Dicer1 knockdown. βIII-tubulin was used as a loading control. **(I)** The optical densities of activated caspase-3 relative to βIII-tubulin from H were normalized to NC siRNA group, and averaged from three independent culture preparation. Student’s *t-test* was used to compare the differences, ***p=0.0008 for comparison in CNs and ***p=0.002 for comparison in HNs.

### Aβ42 oligomer decreased Dicer1 and Nrf2

As above (Figure.1), we found that Dicer1 was reduced in APP/PS1 mice which begin Aβ deposition by six months, with extensive Aβ plaque deposits in the hippocampus and cortex by nine months (Jankowsky et al., 2004). Thus, we investigated the effect of Aβ42 on Dicer1 expression with primary murine cortical neuronal cultures. Initially, we tested the effects of neuronal cultures subject to Aβ42 oligomer treatment, which reduced Dicer1 mRNA in CNs (t(2)=5.879, p=0.0277, paired *t*-test) and HNs (t(2)=4.936, p=0.0387, paired *t*-test) and increased B1 RNA in CNs (t(2)=4.453, p=0.0469, paired *t*-test) and HNs (t(16)=6.177, p=0.0252, paired *t*-test) (Supplemental Figure. 2). As expected, Aβ42 treatment reduced Dicer1 compared to sham treatment (p=0.00001),which was rescued by N-acetyl-cysteine, an antioxidant (p=0.032)(F(2,6)=73.55, p=0.00001, *ANOVA*) (Figures.3A and 3B). Simultaneously, we examined total Nrf2 and found that Nrf2 was reduced by Aβ42 (p=0.0005), which was rescued by addition of N-acetyl-cysteine (p=0.0009) (F(2,6)=37.95, p=0.0004, *ANOVA*) (Figures. 3A and 3C). Since Keap1 regulates Nrf2 stability, we examined Keap1 protein levels and found that Aβ42 increased Keap1(p=0.0014), which was reduced by addition of N-acetyl-cysteine (p=0.0003) (F(2,6)=42.78, p=0.0003, *ANOVA*) (Figures. 3A and 3D). Since Nrf2 affects anti-oxidant gene transcription via binding with AREs in nucleus (Itoh et al., 2004), we further examined the levels of nucleic or cytosolic Nrf2 under Aβ42 treatment or those under simultaneous addition of N-acetyl-cysteine. Aβ42 reduced either nucleic Nrf2 (p=0.0326) (Figures. 3E and 3G) or cytosolic Nrf2 (p=0.0076) (Figures. 3F and 3H), which was rescued by addition of N-acetyl-cysteine (p=0.00001 for nucleic)(F(2,6)=73.82, p=0.00001, *ANOVA*) or (p=0.0419 for cytosolic)(F(2,6)=11.73, p=0.0084, *ANOVA*), respectively (Figures. 3G and 3H).

**Figure 3.**
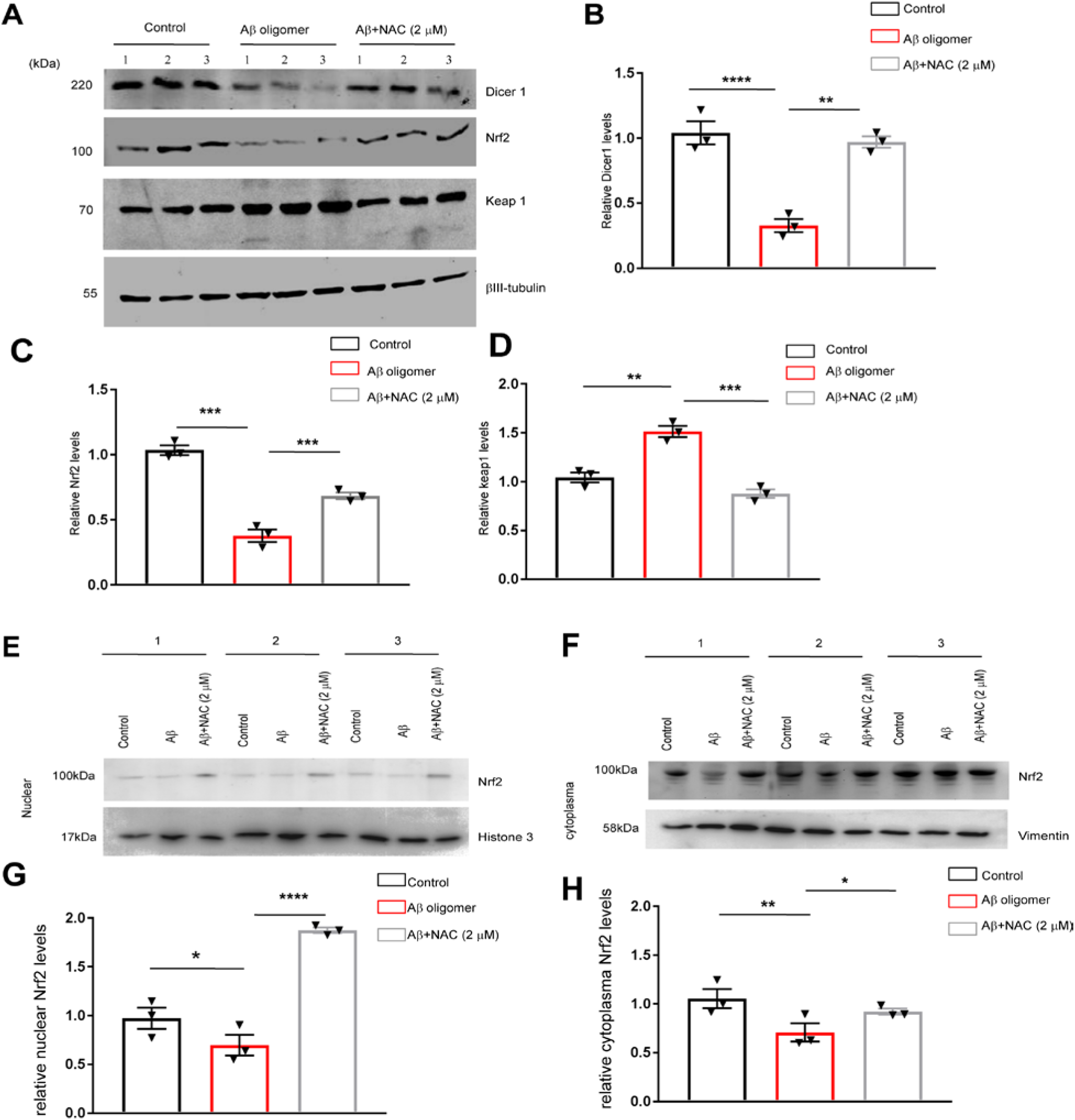
Aβ42 oligomer reduced Dicer1 which was coupled with Nrf2 reduction, the effects rescued by NAC. The primary mouse cortical neurons were treated with Aβ42 oligomer (100 nM) and antioxidant NAC (2 μM) for 48 h. **(A)** The neuronal cultures were harvested after the treatment, homogenized, and subject to centrifugation at 12,000 X g. The supernatants were collected for western blot against Dicer 1(1:600), Nrf2 (1:600), and Keap1(1:600). βIII-tubulin (1:1000) was used as an internal loading control. **(B)** The optical densities of Dicer1/βIII-tubulin were normalized to sham treatment and averaged from three independent culture preparation in A. One-way ANOVA followed by Tukey’s *post hoc* test was used to compare the differences. Control *vs*. Aβ42 oligomer treatment, ****p=0.00001; Aβ42 oligomer *vs*. Aβ42 oligomer plus NAC treatment (Aβ+NAC), *p=0.032. **(C)** The optical densities of Nrf2/βIII-tubulin were normalized to sham treatment and averaged from three independent culture preparation in A. One-way ANOVA followed by Tukey’s *post hoc* test was used to compare the differences. Control *vs*. Aβ42 oligomer treatment, ***p=0.0005; Aβ42 oligomer *vs*. Aβ+NAC, ***p=0.0009. **(D)** The optical densities of Keap1/βIII-tubulin were normalized to sham treatment and averaged from three independent culture preparation in A. Control *vs*. Aβ42 oligomer, **p=0.0014; Aβ42 oligomer *vs*. Aβ+NAC, ***p=0.0003. The neurons were treated with Aβ42 oligomer (100 nM) and antioxidant NAC (2 μM) for 24 h. **(E)** The nucleic proteins were extracted and blotted against Nrf2 (1:600), and histone3 (1:800) served as a loading control; **(F)** the cytoplasmic proteins were isolated and blotted against Nrf2 (1:600), and vimentin (1:500) served as a loading control. **(G)** The optical densities of nucleic Nrf2 relative to histone 3 averaged from three independent culture preparation in E. One-way ANOVA followed by Tukey’s *post hoc* test was used to compare the differences. Control *vs*. Aβ42, *p=0.0326; Aβ42 *vs*. Aβ+NAC,****p=0.00001. **(H)** The optical densities of cytosolic Nrf2/vimentin were normalized to sham treatment and averaged from three independent culture preparation in F. One-way ANOVA followed by Tukey’s *post hoc* test was used to compare the differences. Control *vs*. Aβ42 oligomer, *p=0.0076; Aβ42 oligomer *vs*. Aβ+NAC, *p=0.0419.

### Nrf2 regulated Dicer1 expression

Nrf2 is a master regulator of transcription of anti-oxidant genes. We hypothesized that Nrf2 may regulate Dicer1 expression. To reach the purpose, we initially tested Dicer1 expression in SK-N-BE(2) cells with overexpression of Nrf2 compared to the cells transfected with vehicle plasmid (p=0.0017) (F(2,6)=30.82, p=0.0007, *ANOVA*)(Figures. 4A and 4C). Indeed, overexpression of Nrf2 significantly increased Dicer1 expression when compared to the cells transfected with vehicle plasmid (p=0.0021)(F(2,6)=21.96, p=0.0017, *ANOVA*)(Figures. 4A and 4B). To further consolidate the regulation of Dicer1 by Nrf2, we further examined Dicer1 expression in Neuro-2a cells with overexpression of Nrf2 (t(2)=6.473, p=0.024, paired t-test) (Figures.4D and 4F), which significantly increased Dicer1 expression when compared to the cells transfected with vehicle plasmid (t(2)=10.86, p=0.0004, paired *t*-test)(Figures. 4D and 4E). We further scanned the promoter of Dicer1 with software (promoter 2.0 prediction server) and found three AREs motif in the proximal promoter region around transcription start site (tss). We cloned the ARE1-ARE3 into PGL6-basic plasmid containing firefly luciferase-coding sequence (-Luc), respectively, and also cloned the promoters containing different ARE motif into PGL6-basic plasmid, respectively (F(17,54)=27.87, p=0.00001, *ANOVA*) (Figure. 4G). Transfection of Nrf2-expressing plasmid (pCMV Nrf2) increased the ratios between *firefly* and *renilla* in constructs containing ARE1(p=0.000219) or ARE3 motif (p=0.043) when compared to the cells transfected with vehicle plasmid, pCMV3. When compared the constructs containing different AREs under transfection with pCMV Nrf2, we found that the ratios of firefly/renilla from ARE1 constructs were significantly from the construct ARE2 (p=0.000825), ARE3 (p=0.0093), and PGL6-basic (p=0.00041), respectively. Transfection of pCMV Nrf2 increased firefly/renilla in promoter 1(p=0.00047), promoter 2 (p=0.000135), promoter 3 (p=0.000114), and mutant promoter 1 (p=0.048) when compared to the cells transfected with pCMV3. When compared different promoter constructs under transfection with pCMV Nrf2, we found that the ratios of firefly/renilla from promoter1 were significantly from promoter 0 (p=0.000361) and mutant promoter1(p=0.009) but not different with promoter 2 (p=0.194) or promoter 3 (p=0.254). Basing on these data, we projected that ARE1 motif was critical for Nrf2 regulation of *Dicer1* gene transcription and expression.

**Figure 4.**
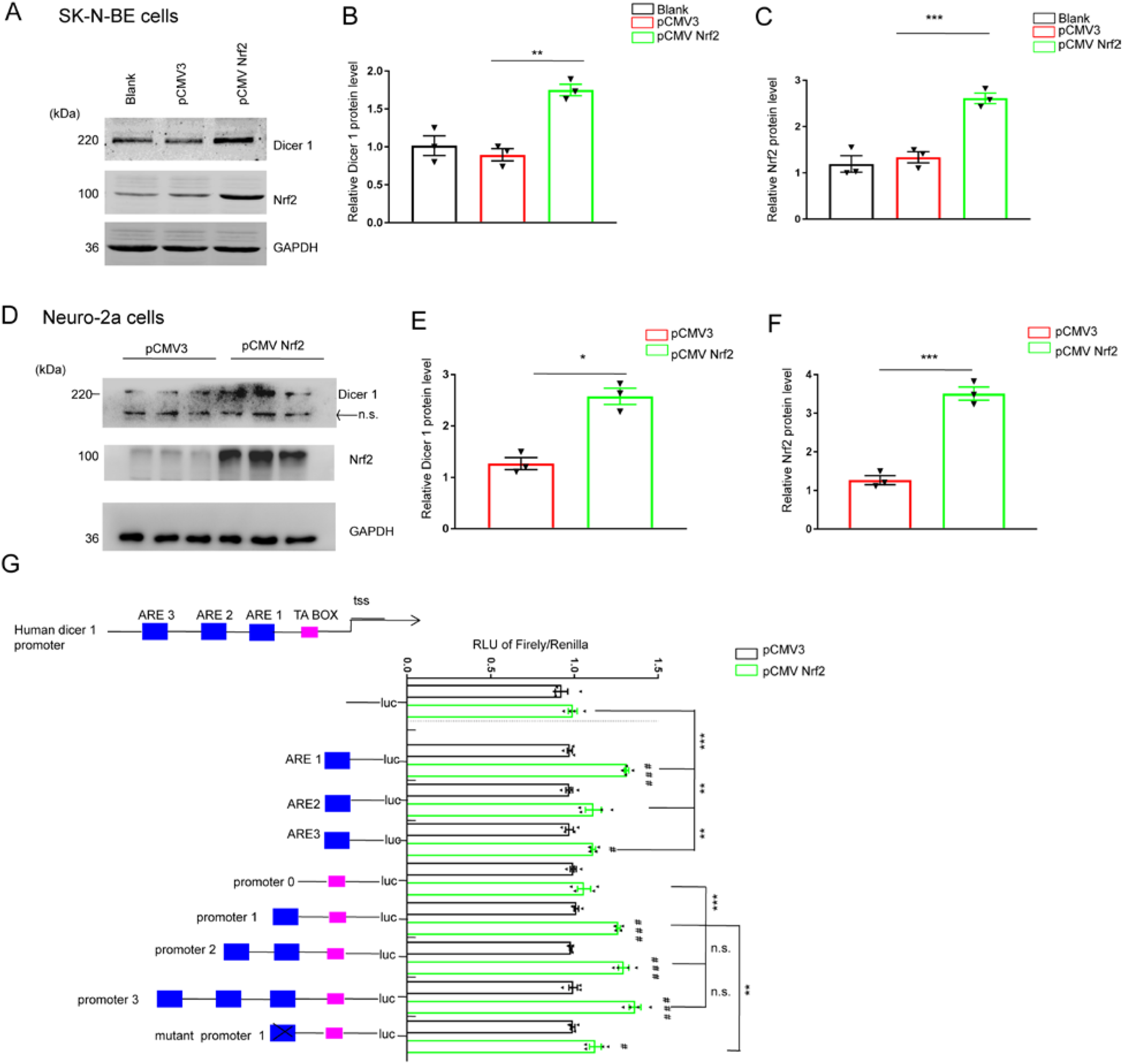
Nrf2 regulated Dicer1 production via Nrf2-ARE pathway. **(A)** SK-N-BE(2) cells were left untreated (Blank) or transfected with a human Nrf2-expressing plasmid (pCMV Nrf2, 1.5 μg) or equal amount of vehicle plasmid (pCMV3) in 6-well plates by use of lipofectamine 2000 in DMEM plus 10% FBS. Forty-eight hours after transfection, the cells were harvested for western blot against Nrf2 and Dicer1. GAPDH was used as a loading control. **(B)** The protein levels of Dicer1 relative to GAPDH were normalized to pCMV group and averaged from three independent culture preparation in A. One-way ANOVA followed by Tukey’s *post hoc* test was used to compare the differences, pCMV3 *vs*. pCMV3 Nrf2, **p=0.0021. **(C)** The protein levels of Nrf2/ GAPDH were normalized to pCMV group and averaged from three independent culture preparations in A. One-way ANOVA, followed by Tukey’s *post hoc* test was used to compare the difference, pCMV3 *vs*. pCMV3 Nrf2, **p=0.0017. **(D)** Neuro-2a cells transfected with a murine Nrf2-expressing (pCMV Nrf2) or vehicle plasmid (pCMV3) by use of lipofectamine 2000 for 48 h. At the end of the culture, the cells were harvested for western blot against Nrf2 and Dicer1. GAPDH was used as a loading control. **(E)** The protein levels of Dicer1/ GAPDH were normalized to pCMV3 group and averaged from three independent culture preparation in D. Student’s *t-test* was used to compare the differences, pCMV3 *vs*. pCMV3 Nrf2, *p=0.0004. **(F)** The protein levels of Nrf2/GAPDH were normalized to pCMV3 group and averaged from three independent culture preparation in D. Student’s *t-test* was used to compare the differences, pCMV3 *vs*. pCMV3 Nrf2, ***p=0.024. **(H)** The upper panel showed scheme of Dicer 1 gene promoter containing three predicted ARE sites (blue square), TA box (pink red), tss indicating transcriptional start site (tss). HEK293 T cells were seeded at a density of 3 X10^4^ cells per well in 48-well plates. At ∼70% confluency, the cells were transfected with individual construct (150 ng), *Renilla* luciferase plasmid (7.5 ng), and PCMV3 or pCMV Nrf2 plasmid expressing human Nrf2 (150 ng) by use of lipofectamine 2000 for 48 h. At the end of culture, the cells were harvested and subject to luciferase activity assay. The results were averaged from five independent culture preparation. One-way ANOVA followed by Tukey’s *post hoc* test was used to compare the differences. Comparing co-transfection of pCMV3 Nrf2 with co-transfection of pCMV3 in individual construct, ^###^ p=0.000219 in ARE1, ^#^p=0.043 in ARE3, ^###^p=0.00047 in promoter 1, ^###^p=0.000135 in promoter2, ^###^p=0.000114 in promoter3, and ^#^p=0.048 in mutant promoter 1. Under the condition of co-transfection of pCMV3 Nrf2, ARE1 *vs*. pGL6-luc, ***p=0.00041; ARE1 *vs*. ARE2, **p=0.000825; ARE1 *vs*. ARE3, **p=0.0093; promoter 1 *vs*. promoter 0, ***p=0.000361; promoter 1 *vs*. mutant promoter 1, **p=0.009, promoter 1 *vs*. promoter 2, p=0.194 (n.s); promoter 1 *vs*. promoter 3, p=0.254 (n.s).

### Overexpression of Dicer1 recovered neurotoxicity by Aβ42 oligomer

Since we have demonstrated that Dicer1 depletion compromised cell survival (Figure 2), we wondered whether overexpression of Dicer1 mitigated Aβ42-induced neurotoxicity. Aβ42 oligomer treatment reduced the protein levels of Dicer1 (p=0.033) compared to sham treatment, which was rescued by infection with Ad-Dicer1-T2A:EGFP virus compared to infection with vehicle virus, Ad-EGFP (p=0.0074) (F(2,6)=3.15, p=0.0213, *ANOVA*) (Figures. 5A and 5B). Aβ42 oligomer treatment increased activated caspase 3 compared to sham treatment (p=0.0061), which was rescued by infection with Ad-Dicer1-T2A:EGFP virus compared to infection with Ad-EGFP (p=0.023) (F(2,6)=2.05, p=0.0438, *ANOVA*) (Figures. 5A and 5C). Under expectation, Aβ42 oligomer decreased neuronal survival (p=0.00067), which was significantly rescued by infection with Ad-Dicer1-T2A:EGFP virus (p=0.047)(F(2,6)=2.46, p=0.0362, *ANOVA*) (Figure. 5D). Notably, Aβ42 oligomer increased the secretion of IL-1β (Figure. 5E) (p=0.023) or IL-18 (Figure. 5F) (p=0.019), which was decreased by infection with Ad-Dicer1-T2A:EGFP virus (F(2,6)=3.18, p=0.0263, *ANOVA*) (Figure. 5E, p=0.005 for IL-1β secretion) and (F(2,6)=2.72, p=0.0328, *ANOVA*)(Figure. 5F, p=0.04 for IL-18 secretion). Dicer1 knockdown disrupted neurite in primary cortical neurons (t(16)=2.51, p=0.011, paired *t*-test) (Supplemental Fig3), an early pathological change in AD. Similarly, Aβ42 oligomer treatment trimmed neurite (p=0.025), which was significantly recovered by overexpression of Ad-Dicer1-T2A:EGFP virus (p=0.047) (F(2, 447)=3.24, p=0.0285, *ANOVA*) (Figures. 5G and 5H).

**Figure 5.**
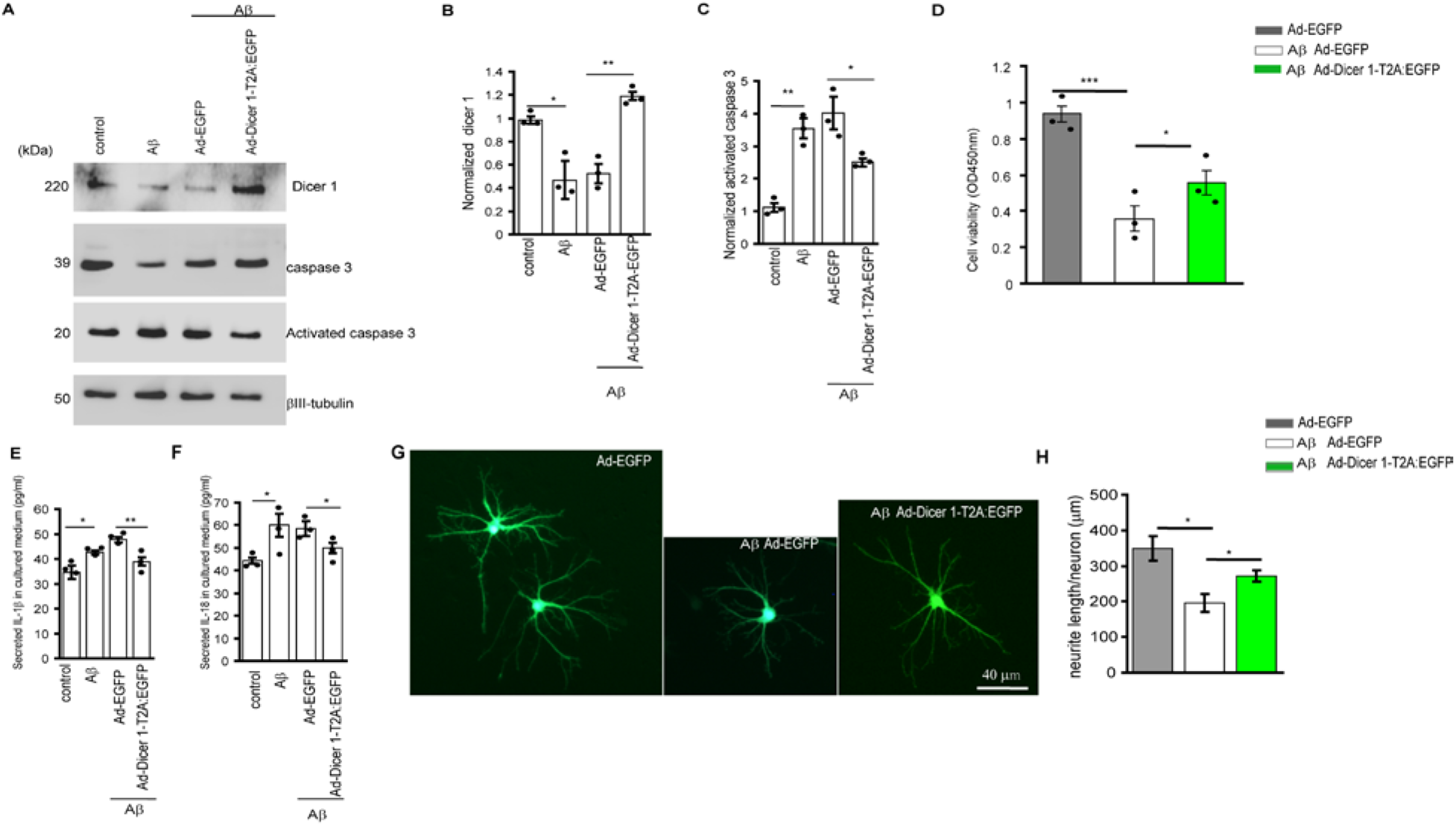
Overexpression of Dicer 1 reduced Aβ42 oligomer-mediated secretion of IL-1β and IL-18, apoptosis, and neurite deficit in primary hippocampal neuronal cultures. (A) The primary murine hippocampal neurons at DIV 3 were treated with Aβ42 oligomer (100 nM) for 24 h or subject to sham treatment in neurobasal medium plus 2% B27. In a parallel experiment, the medium was replaced with fresh neurobasal medium plus 2% B27 following Aβ42 treatment, and then infected with vehicle virus, Ad-EGFP or Ad-dicer1-T2A:EGFP virus (5 X 10^7^ vg/mL) for 48 h. At the end of treatment, the cultures were harvested and homogenized for western blot against Dicer1, caspase 3, and activated caspase 3. βIII-tubulin was used as a loading control. **(B)** Dicer1 relative to βIII-tubulin were normalized to sham treatment and averaged from three independent culture preparation. One-way ANOVA followed by Tukey’s *post hoc* test was used to compare the differences. Dicer1 was reduced by Aβ42 oligomer treatment compared to sham treatment (*p=0.033) and the reduction of Dicer1 by Aβ42 was rescued by infection of Ad-dicer1-T2A:EGFP compared to infection by vehicle virus (**p=0.0074). **(C)** Activated caspase3 relative to βIII-tubulin were normalized to sham treatment and averaged from three independent culture preparation. One-way ANOVA followed by Tukey’s *post hoc* test was used to compare the differences. Activated caspase3 was increased by Aβ42 oligomer compared to sham treatment (**p=0.0061), which was reduced by infection of Ad-dicer1-T2A:EGFP compared to infection by vehicle virus (*p=0.023). **(D)** The loss of cell viability of hippocampal neurons treated by Aβ (***p=0.00067) was significantly rescued by infection with Ad-dicer1-T2A:EGFP compared to infection by vehicle virus (*p=0.047). The values were averaged from quadruplicate cultures with three independent culture preparation. The cultured media were also collected from the cultures treated as in A and subject to ELISA detection of IL-1β and IL-18. **(E)** Aβ42 oligomer treatment increased IL-1β in the supernatants (*p=0.023) which was reduced by infection of Ad-dicer1-T2A:EGFP compared to infection by vehicle virus (**p=0.005). The values were averaged from quadruplicate cultures with three independent cell preparation and One-way ANOVA followed by Tukey’s *post hoc* test was used to compare the differences. **(F)** The culture media were collected and subject to ELISA detection of IL-18 as in E. Aβ treatment increased IL-8 (*p=0.019) which was reduced by infection of Ad-dicer1-T2A:EGFP compared to infection by empty virus (*p=0.04). The values were averaged from quadruplicate cultures with three independent cell preparation and One-way ANOVA followed by Tukey’s post hoc test was used to compare the difference. **(G)**Representative hippocampal neurons (150 neurons for each type of treatment) infected by Ad-EGFP virus (Ad-EGFP), Aβ42 treatment followed by infection with Ad-EGFP virus (Aβ Ad-EGFP) and Aβ42 treatment followed by infection with Ad-Dicer1-T2A:EGFP virus (Aβ Ad-Dicer1:T2A:EGFP) were indicated. **(H)** Neurite lengths of hippocampal neurons with treatment from G were calculated and averaged. Aβ42 treatment reduced neurite length compared to sham treatment (*p=0.025) which was rescued by infection with Ad-dicer 1-T2A:EGFP virus (*p=0.047). The neurite length averaged from 150 neurons in each treatment and One-way ANOVA followed by Tukey’s *post hoc* test was used to compare the differences.

### Overexpression of Dicer1 improved spatial learning in APP/PS1 mice

We further explored the effect of overexpression of Dicer1 in APP/PS1mice. Consistent with previous observation (Figure. 1A), Dicer1 staining was decreased in CA3 hippocampus compared to WT littermate at the age of 3.5 months; injection of Ad-Dicer1:T2A:EGFP significantly increased Dicer1 staining in CA3 hippocampus in APP/PS1 mice (Figure. 6A). CA3 hippocampi of APP/PS1mice contained stronger staining for activated caspase 3 compared to those of WT littermate, which was reduced by injection of Ad-Dicer1:T2A:EGFP virus (Figure. 6B). In parallel experiment, we isolated CA3 hippocampi and subjected the homogenates to western blot against Dicer1 and caspase 3. Consistent with the immunofluorescence, there was lower level of Dicer1 in APP/ PS1mice than WT (p=0.0217) which was recovered by injection of Ad-Dicer1:T2A:EGFP (p=0.0093) (p=0.00236, Kruskal-Wallis test) (Figure. 6D); there was higher level of activated caspase 3 in APP/PS1mice than WT (p=0.00681) which was reduced by injection of Ad-Dicer1:T2A:EGFP (p=0.0067) (p=0.0036, Kruskal-Wallis test) (Figure. 6E). We further investigated whether overexpression of Dicer1 had effect on spatial learning and thus examined behavior with Morris Water Maze. During the training period on days 1-6 after virus injection, the latency to reach platform for APP/PS1 mice was significantly reversed by injection with Ad-Dicer1:T2A:EGFP virus compared to injection with Ad1-EGFP virus (p=0.008) on day 5 (F (2, 42) =9.042 p=0.0057, *ANOVA*), and (p=0.00001) on day 6 (F (2, 42) =13.12, p=0.0025, ANOVA) (Figure. 7A). When combining timepoint and treatment, the latency was significantly reversed by injection with Ad1-Dicer1:T2A:EGFP virus (p=0.0001) (Fday (5, 174) = 36.44, p=0.00001; Ftreatment (2, 174) = 2.045, p=0.0241; Finteraction (75, 174) = 2.3, p=0.0148, two-way *ANOVA*) (Figure. 7A). On day 7 after a 60-s probe test for searching platform, we monitored animal’s swimming pattern, distance, speed, and the amount of time spent in each of the four quadrants. Without difference on swimming distance (F (2, 42)=1.19, p=0.3141, *ANOVA*) (Figure. 7D) and swimming speed (F (2, 42) = 1.49, p=0.0948, *ANOVA*) (Figure. 7E), APP/PS1mice spent less time in target quadrant than WT mice (p=0.0023), which was reversed by injection with Ad-Dicer1:T2A:EGFP virus (p=0.0087) (F(2, 42)=5.43 P=0.0092; F_quadrant_ (3, 116) = 4.576, p=0.1122; F_treatment_ (2,116) = 0.1032, p=1; F_interaction_ (45, 116) = 1.334. p=0.0046, two-way ANOVA) (Figure. 7B). Similarly, the AD mice crossed the platform less frequently than WT mice (p=0.039), but the chance was increased by injection with Ad-Dicer1:T2A:EGFP virus (p=0.043) (F(2, 42)= 4.967, p=0.0116, *ANOVA*) (Figure. 7C).

**Figure 6.**
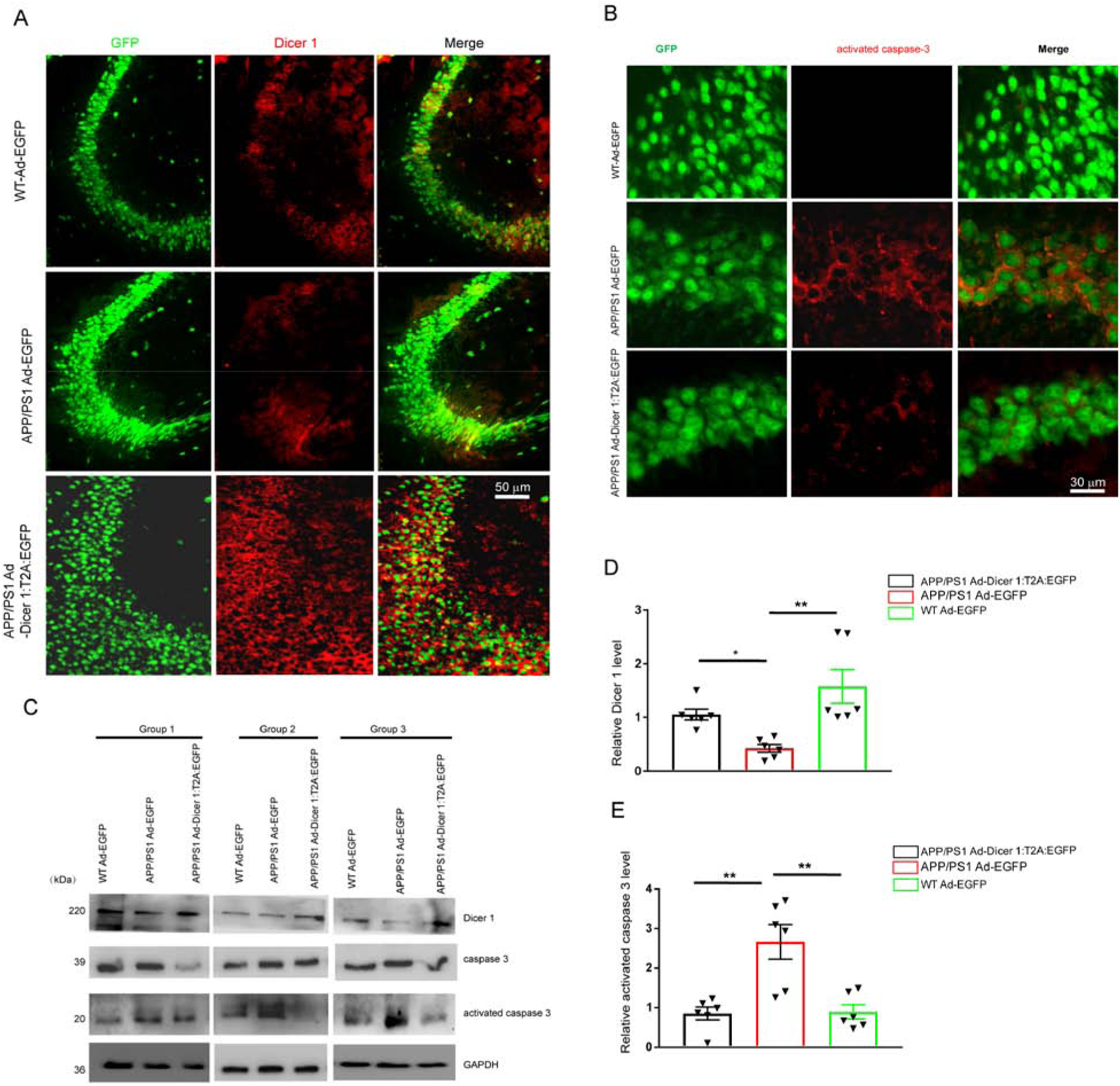
Overexpression of Dicer1 in APP/PS1hippocampus inhibited activated caspase 3. APP/PS1 mice or WT littermate at the age of 3.5-/4-month were injected with Ad-EGFP or Ad-Dicer1-T2A:EGFP virus (1.2 X10^9^ vg/mL) in CA3 hippocampi. 17 days after injection, the injected mice were perfused and each brain was sagittally cut into ∼60 sections at 10 μm-thick for each section. The sections every other six sections were chosen for double staining of Dicer1 (red) (**A**) and activated caspase 3 (red) (**B**). The typical images of CA3 hippocampal region were indicated. **(A)** Dicer1 staining (red) in APP/PS1 CA3 hippocampus was reduced compared to those in WT littermate and infection with Ad-Dicer 1-T2A:EGFP virus dramatically increased Dicer1 staining in CA3 hippocampus. The images were typical of 60 sections from six mice. Bar, 50 μm. **(B)** The staining of activated caspase 3 (red) in APP/PS1mice was increased compared to those in WT and this increase in staining was mitigated by infection with Ad-Dicer1-T2A:EGFP virus. The images were typical of 60 sections from six mice. Bar, 30 μm. **(C)** The CA3 hippocampi were harvested, homogenized, and subjected to centrifugation at 12,000 X g, and the supernatants were used for western blot against Dicer1, caspase3, and activated caspase3, and typical blots were indicated. (**D**) Dicer1 was reduced in APP/PS1 hippocampi compared to WT (*p=0.0217), which was increased by infection with Ad-Dicer1-T2A:EGFP virus (**p=0.0093). The optical densities averaged from six mice in each group and Kruskal-Wallis test was used to compare the differences. (**E**) Activated caspase 3 relative to GAPDH was increased in APP/PS1 hippocampi compared to WT (**p=0.00681), which was reduced by infection with Ad-Dicer1-T2A:EGFP virus (**p=0.0067). The optical densities averaged from six mice in each group and Kruskal-Wallis test was used to compare the differences.

**Figure 7.**
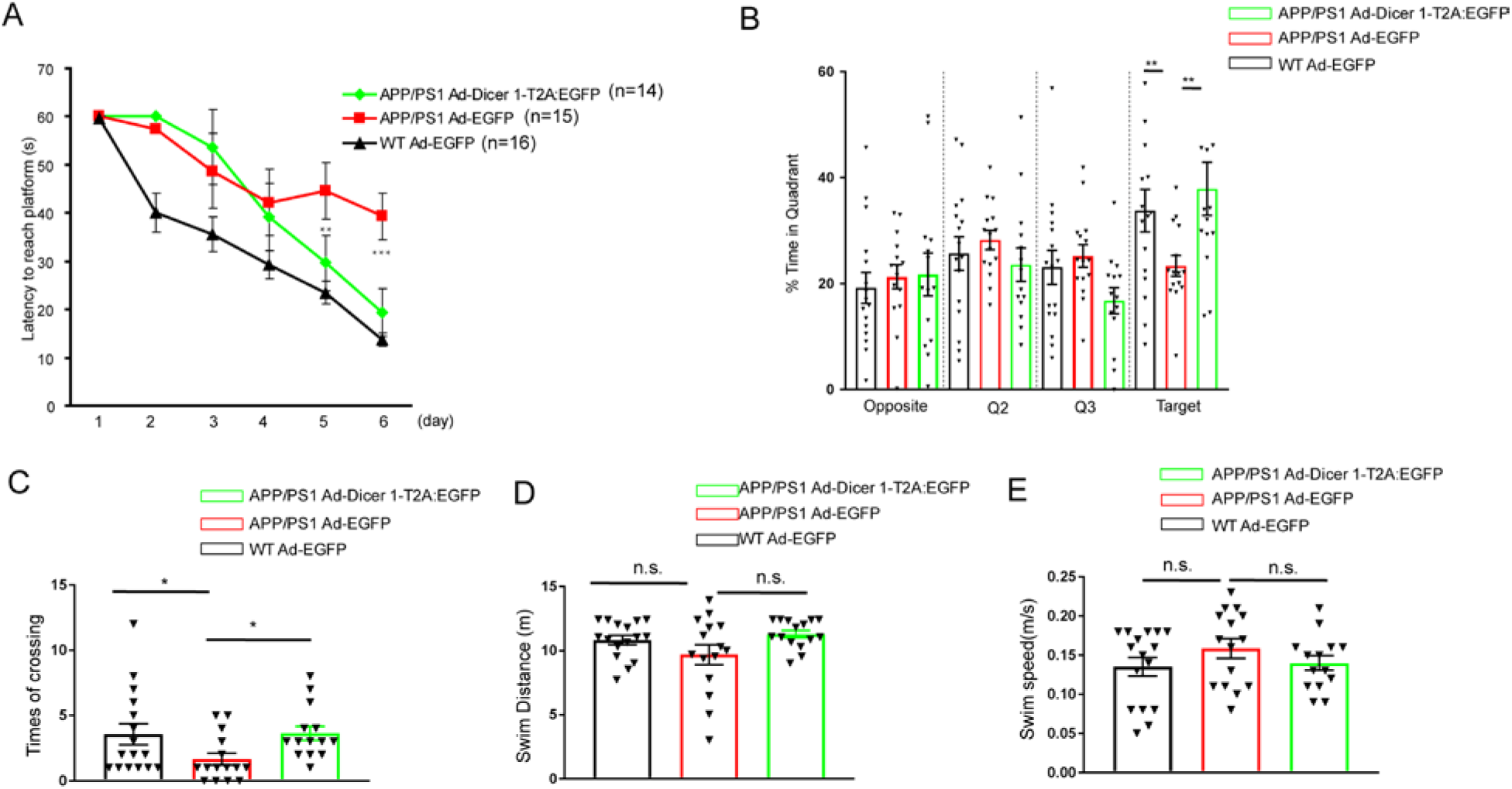
Overexpression of Dicer 1 in CA3 hippocampus significantly improved spatial learning in APP/PS1 mice. Ad-Dicer1-T2A:EGFP virus (1.2 X 10^9^ vg/mL) or equal amount of Ad-EGFP virus were injected into CA3 hippocampus in APP/PS1or WT mice and behavior test was conducted at 17-23 days after injection. **(A)** Learning curve of the mice from WT mice injected with Ad-EGFP virus (n=16), APP/PS1 mice injected with Ad-EGFP virus (n=15) or Ad-Dicer1-T2A:EGFP virus (n=14). The differences of latency to reach platform were analyzed by use of two-way ANOVA with repeated measures against day and treatment (virus injected), p_day_ =0.00001, p_treatment_=0.0241, p_interaction_=0.0148. Tukey’s *post hoc* test was further used to compare the differences between APP/PS1 mice injected with Ad-Dicer1-T2A:EGFP virus and APP/PS1mice injected with Ad-EGFP virus, **p=0.008 on day 5 and ***p=0.00001 on day 6. **(B)** Preference of APP/PS1mice for the target quadrant was enhanced by Ad-Dicer 1-T2A:EGFP virus injection in CA3 hippocampi. Two-way ANOVA with repeated measures against quadrant and treatment was used to compare the differences, p_quadrant_ =0.1122, p_type_=1, p_interaction_=0.0046. Tukey’s *post hoc* test was further used to compare the differences among groups, APP/PS1 Ad-Dicer 1-T2A:EGFP (n=16) *vs*. APP/PS1 Ad-EGFP(n=15), **p=0.0087; APP/PS1 Ad-EGFP (n=15) *vs*. WT Ad-EGFP (n=16), **p=0.0023. **(C)** Frequencies of crossing target platform was increased by Dicer1 overexpression. Tukey’s *post hoc* test was further used to compare the differences among groups, APP/PS1 Ad-Dicer 1-T2A:EGFP (n=14) *vs*. APP/PS1 Ad-EGFP (n=15); *p=0.043; APP/PS1 Ad-EGFP (n=15) *vs*. WT Ad-EGFP (n=16), *p=0.039. **(D)** Swim distance and **(E)** Swim speed did not show differences among groups (n.s).

## Discussion

In this study, we indicated that Dicer1 was reduced in the hippocampus and cortex of APP/PS1 mice *in vivo* and Dicer1 was reduced by Aβ42 oligomer *in vitro*. Dicer1 depletion led to neurite deficit and apoptosis via oxidative injury-mediated inflammasome activation such as increased ROS production and activated caspase3, damaged mitochondrial transmembrane potential, and IL-1β and IL-18 secretion. We further indicated that Nrf2 increased Dicer1 expression which was linked with Keap1-Nrf2-ARE signaling pathway. Overexpression of Dicer1 increased neuronal survival and recovered neurite deficit by Aβ42 oligomer *in vitro*, and *in vivo* Dicer1 overexpression in CA3 hippocampus improved spatial learning in APP/PS1 mice. Altogether, we reveal a novel role of Dicer1 in AD and a novel regulation of Dicer1. In our study, reduction of Dicer1 was detected as early as around 4 months in APP/PS1 mice which begin Aβ deposition at the age of 6 months (Jankowsky et al., 2004). Furthermore, knockdown of Dicer1 in cultured neurons induced neurite deficit before soma degeneration, and intrahippocampal injection of Dicer1-expressing virus improved spatial learning. These evidence suggest that reduction of Dicer1 is an early event in AD and targeting Dicer1 may provide a new strategy for AD therapy with emphasis on the early stage of this disorder.

Oxidative stress is thought to be central in the pathogenesis of AD, for example, oxidative stress exacerbates insulin resistance in AD brain (De Felice et al., 2014) and disrupts mitochondria (Lin and Beal, 2006), resulting in failure to produce sufficient ATP for Na^+^-K^+^-ATPase and Na^+^-Ca^++^ exchanger to maintain ion homeostasis in cells (Reeves et al., 1986; Gloor, 1997; Kip and Strehler, 2007). Dicer1 depletion may be an internal sources contributing to oxidative stress by accumulation of Alu RNA (Tarallo et al., 2012). Intrahippocampal injection of Nrf2-expressing lentivirus improves spatial learning in APP/PS1 mice (Kanninen et al., 2009) but Nrf2 promotes cancer cell proliferation(Jaramillo and Zhang, 2013). As such, manipulation of Nrf2 expression may lead to undesired side effects although it indicates potential benefit in AD brain (Kanninen et al., 2009). By contrast, appropriate expression of Dicer1 may be an alternative strategy to oppose oxidative stress in AD brain. Dicer1 is ubiquitously expressed in the brain including in neurons, astroglia, and oligodendrocytes (Cuellar et al., 2008; Shin et al., 2009; Tao et al., 2011). Thus, Dicer1 reduction in APP/PS1 brain and secondary neurodegeneration are the combinatory effects from neurons and glia. Therefore, it is important to determine the contribution of Dicer1 reduction in cell type in AD brain for therapy purpose in the future. Nrf2 mediates Dicer1 expression via Keap1-Nrf2-ARE signaling pathway, and decreased Nrf2-ARE signaling may mediate the reduction of Dicer1 in AD brain as projected from our study. Overexpression of Dicer1 may serve as a forward regulator of Nrf2, which is unclear yet.

Dicer1 is central to the processing from pre-microRNAs to mature microRNAs, which regulate neuronal survival and neuritogenesis (Im and Kenny, 2012; Carrella et al., 2015). Deletion of functional Dicer1 has been found to induce accumulation of Alu RNA and activate NLRP3 inflammasome and oxidative damages in age-related macular degeneration(Kaneko et al., 2011; Tarallo et al., 2012). Conditional ablation of Dicer1 in oligodendrocytes induces neuronal impairment *via* demyelination, oxidative damage, inflammatory astrocytosis and microgliosis in mice (Shin et al., 2009). There is also evidence showing that conditional knockout of Dicer1 in dopaminergic neurons of ventral midbrain leads to neuronal loss through inhibiting micoRNAs biosynthesis(Chmielarz et al., 2017). Similarly, blocking microRNAs biosynthesis by knocking out Dicer1 disrupts morphogenesis in the cortex and hippocampus (Davis et al., 2008). Apart from Alu RNA accumulation, microRNAs deficiency due to Dicer1 depletion may play unexpected roles in AD neurodegeneration as Alu RNA, which is of great interest to explore.

In summary, we revealed novel roles of Dicer1 in AD brain and a novel regulation of Dicer1. Our findings suggest Dicer1 may be a target in AD therapy.

## Supporting information

supplemental results

## Acknowledgements

This study is supported by Natural Science Foundation of Zhejiang Province (LY18H120003) and by funding (QTJ10001) from Wenzhou Medical University.

## Author contribution

YW designed, conducted the experiments, analyzed data, and wrote the first draft; MLL conducted the experiments; JZ began the characterization of Dicer1 expression in APP/PS1 mice; SZW conceived of this project, analyzed data, and wrote the final manuscript.

## References

Belaidi AA, Bush AI (2016) Iron neurochemistry in Alzheimer’s disease and Parkinson’s disease: targets for therapeutics. J Neurochem 139 Suppl 1:179–197.

Bernstein E, Caudy AA, Hammond SM, Hannon GJ (2001) Role for a bidentate ribonuclease in the initiation step of RNA interference. Nature 409:363–366.

Carrella S, D’Agostino Y, Barbato S, Huber-Reggi SP, Salierno FG, Manfredi A, Neuhauss SC, Banfi S, Conte I (2015) miR-181a/b control the assembly of visual circuitry by regulating retinal axon specification and growth. Developmental neurobiology 75:1252–1267.

Chen ZH, Saito Y, Yoshida Y, Sekine A, Noguchi N, Niki E (2005) 4-Hydroxynonenal induces adaptive response and enhances PC12 cell tolerance primarily through induction of thioredoxin reductase 1 via activation of Nrf2. J Biol Chem 280:41921–41927.

Chmielarz P, Konovalova J, Najam SS, Alter H, Piepponen TP, Erfle H, Sonntag KC, Schutz G, Vinnikov IA, Domanskyi A (2017) Dicer and microRNAs protect adult dopamine neurons. Cell Death Dis 8:e2813.

Cuellar TL, Davis TH, Nelson PT, Loeb GB, Harfe BD, Ullian E, McManus MT (2008) Dicer loss in striatal neurons produces behavioral and neuroanatomical phenotypes in the absence of neurodegeneration. Proc Natl Acad Sci U S A 105:5614–5619.

Davis TH, Cuellar TL, Koch SM, Barker AJ, Harfe BD, McManus MT, Ullian EM (2008) Conditional loss of Dicer disrupts cellular and tissue morphogenesis in the cortex and hippocampus. J Neurosci 28:4322–4330.

De Felice FG, Lourenco MV, Ferreira ST (2014) How does brain insulin resistance develop in Alzheimer’s disease? Alzheimer’s & dementia : the journal of the Alzheimer’s Association 10:S26–32.

Gloor SM (1997) Relevance of Na,K-ATPase to local extracellular potassium homeostasis and modulation of synaptic transmission. FEBS Lett 412:1–4.

Halliwell B (2006) Oxidative stress and neurodegeneration: where are we now? J Neurochem 97:1634–1658.

Hardy J, Selkoe DJ (2002) The amyloid hypothesis of Alzheimer’s disease: progress and problems on the road to therapeutics. Science 297:353–356.

Heneka MT, Kummer MP, Stutz A, Delekate A, Schwartz S, Vieira-Saecker A, Griep A, Axt D, Remus A, Tzeng TC, Gelpi E, Halle A, Korte M, Latz E, Golenbock DT (2013) NLRP3 is activated in Alzheimer’s disease and contributes to pathology in APP/PS1 mice. Nature 493:674–678.

Im HI, Kenny PJ (2012) MicroRNAs in neuronal function and dysfunction. Trends in neurosciences 35:325–334.

Itoh K, Tong KI, Yamamoto M (2004) Molecular mechanism activating Nrf2-Keap1 pathway in regulation of adaptive response to electrophiles. Free Radic Biol Med 36:1208–1213.

Jankowsky JL, Fadale DJ, Anderson J, Xu GM, Gonzales V, Jenkins NA, Copeland NG, Lee MK, Younkin LH, Wagner SL, Younkin SG, Borchelt DR (2004) Mutant presenilins specifically elevate the levels of the 42 residue beta-amyloid peptide in vivo: evidence for augmentation of a 42-specific gamma secretase. Hum Mol Genet 13:159–170.

Jaramillo MC, Zhang DD (2013) The emerging role of the Nrf2-Keap1 signaling pathway in cancer. Genes Dev 27:2179–2191.

Jiang H, Wu M, Liu Y, Song L, Li S, Wang X, Zhang YF, Fang J, Wu S (2017) Serine racemase deficiency attenuates choroidal neovascularization and reduces nitric oxide and VEGF levels by retinal pigment epithelial cells. J Neurochem 143:375–388.

Kaneko H et al. (2011) DICER1 deficit induces Alu RNA toxicity in age-related macular degeneration. Nature 471:325–330.

Kanninen K, Heikkinen R, Malm T, Rolova T, Kuhmonen S, Leinonen H, Yla-Herttuala S, Tanila H, Levonen AL, Koistinaho M, Koistinaho J (2009) Intrahippocampal injection of a lentiviral vector expressing Nrf2 improves spatial learning in a mouse model of Alzheimer’s disease. Proc Natl Acad Sci U S A 106:16505–16510.

Keller JN, Germeyer A, Begley JG, Mattson MP (1997) 17Beta-estradiol attenuates oxidative impairment of synaptic Na+/K+-ATPase activity, glucose transport, and glutamate transport induced by amyloid beta-peptide and iron. J Neurosci Res 50:522–530.

Kip SN, Strehler EE (2007) Rapid downregulation of NCX and PMCA in hippocampal neurons following H2O2 oxidative stress. Ann N Y Acad Sci 1099:436–439.

Kobayashi A, Kang MI, Okawa H, Ohtsuji M, Zenke Y, Chiba T, Igarashi K, Yamamoto M (2004) Oxidative stress sensor Keap1 functions as an adaptor for Cul3-based E3 ligase to regulate proteasomal degradation of Nrf2. Mol Cell Biol 24:7130–7139.

Lin MT, Beal MF (2006) Mitochondrial dysfunction and oxidative stress in neurodegenerative diseases. Nature 443:787–795.

Mark RJ, Hensley K, Butterfield DA, Mattson MP (1995) Amyloid beta-peptide impairs ion-motive ATPase activities: evidence for a role in loss of neuronal Ca2+ homeostasis and cell death. J Neurosci 15:6239–6249.

Mark RJ, Pang Z, Geddes JW, Uchida K, Mattson MP (1997) Amyloid beta-peptide impairs glucose transport in hippocampal and cortical neurons: involvement of membrane lipid peroxidation. J Neurosci 17:1046–1054.

Markesbery WR (1997) Oxidative stress hypothesis in Alzheimer’s disease. Free Radic Biol Med 23:134–147.

Mattson MP (2004) Pathways towards and away from Alzheimer’s disease. Nature 430:631–639.

McWalter GK, Higgins LG, McLellan LI, Henderson CJ, Song L, Thornalley PJ, Itoh K, Yamamoto M, Hayes JD (2004) Transcription factor Nrf2 is essential for induction of NAD(P)H:quinone oxidoreductase 1, glutathione S-transferases, and glutamate cysteine ligase by broccoli seeds and isothiocyanates. The Journal of nutrition 134:3499S–3506S.

Rada P, Rojo AI, Evrard-Todeschi N, Innamorato NG, Cotte A, Jaworski T, Tobon-Velasco JC, Devijver H, Garcia-Mayoral MF, Van Leuven F, Hayes JD, Bertho G, Cuadrado A (2012) Structural and functional characterization of Nrf2 degradation by the glycogen synthase kinase 3/beta-TrCP axis. Mol Cell Biol 32:3486–3499.

Reeves JP, Bailey CA, Hale CC (1986) Redox modification of sodium-calcium exchange activity in cardiac sarcolemmal vesicles. J Biol Chem 261:4948–4955.

Rushworth JV, Griffiths HH, Watt NT, Hooper NM (2013) Prion protein-mediated toxicity of amyloid-beta oligomers requires lipid rafts and the transmembrane LRP1. J Biol Chem 288:8935–8951.

Shin D, Shin JY, McManus MT, Ptacek LJ, Fu YH (2009) Dicer ablation in oligodendrocytes provokes neuronal impairment in mice. Ann Neurol 66:843–857.

Stockwell BR et al. (2017) Ferroptosis: A Regulated Cell Death Nexus Linking Metabolism, Redox Biology, and Disease. Cell 171:273–285.

Tao J, Wu H, Lin Q, Wei W, Lu XH, Cantle JP, Ao Y, Olsen RW, Yang XW, Mody I, Sofroniew MV, Sun YE (2011) Deletion of astroglial Dicer causes non-cell-autonomous neuronal dysfunction and degeneration. J Neurosci 31:8306–8319.

Tarallo V et al. (2012) DICER1 loss and Alu RNA induce age-related macular degeneration via the NLRP3 inflammasome and MyD88. Cell 149:847–859.

