## supplemental results for "Dicer1 is reduced in APPswe/PSEN1dE9 mice and is regulated by Nrf2"

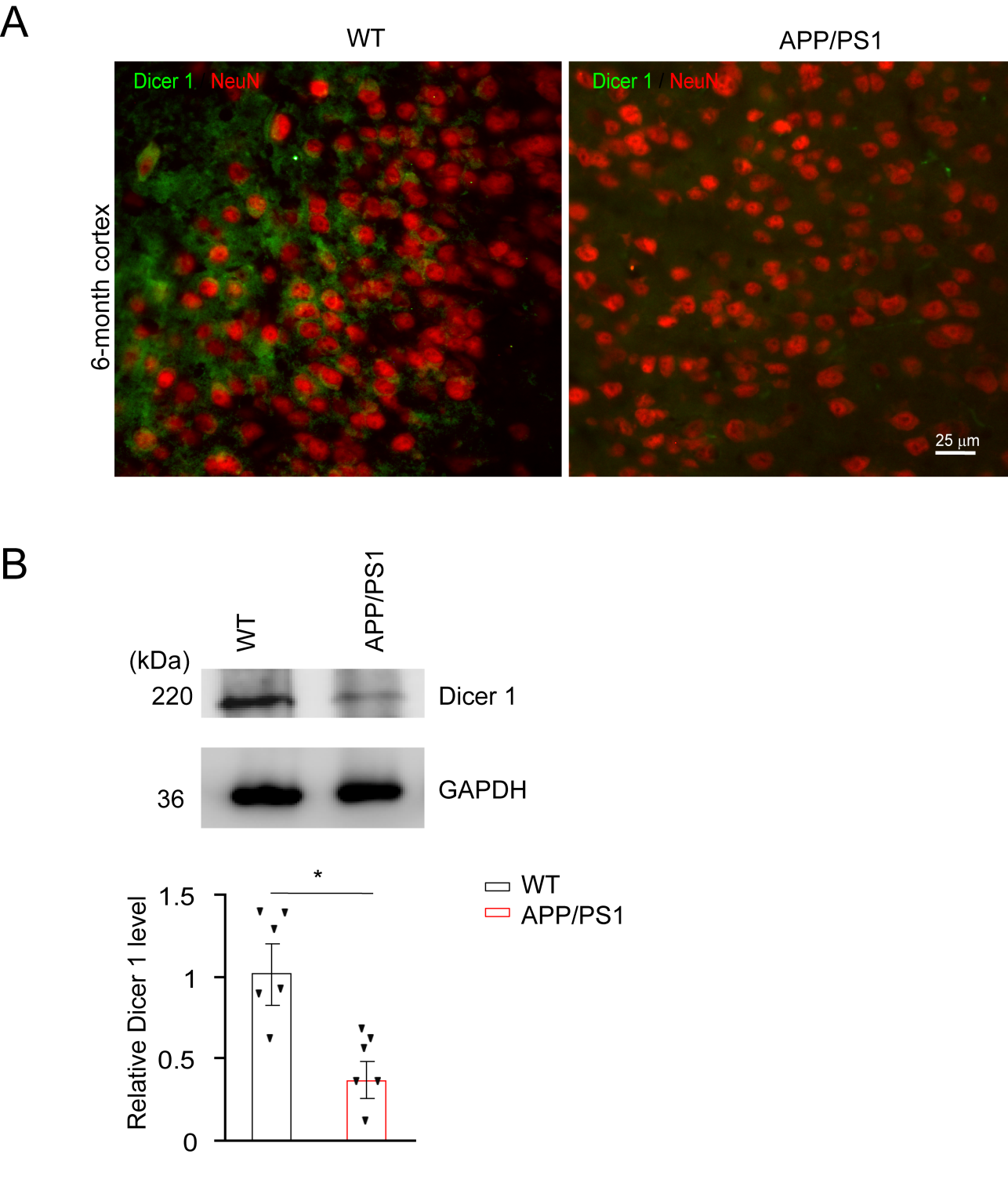

**Figure S1.** **Reduction of Dicer 1 in 6-month-old APP/PS1 cortex.** **(A)** Representative image of Dicer1 staining in parietal cortex with double staining by anti-Dicer1(green) and anti-NeuN(red). (n=3, per genotype). Bar, 25 μm for all panels. **(B)** The homogenates of parietal lobe from WT (n=6) and APP/PS1(n=6) were subject to western blot against Dicer1. GAPDH was used as a loading control. The optical densities of Dicer1 relative to GAPDH averaged from three independent experiments. Mann-Whitney U test was used to compare the differences, *p=0.022.

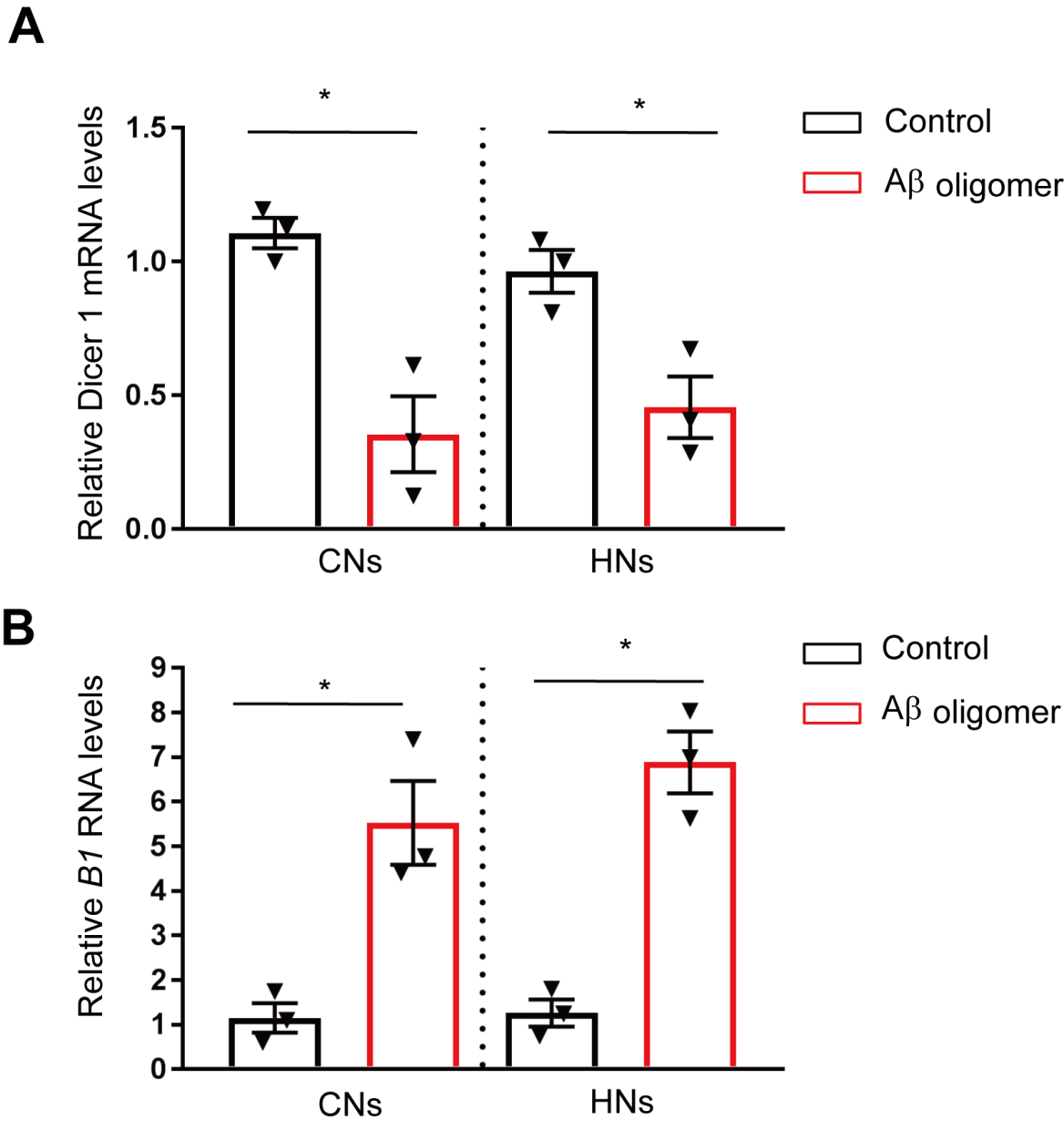

**Figure S2.** **Aβ42 oligomer decreased *Dicer1* mRNA and increased B1 RNA in primary neuronal cultures.** Primary murine cortical neuronal cultures were treated with Aβ42 oligomer (100 nM) for 48 h and subject to real-time quantitative PCR examination of *Dicer1* mRNA (**A**) and B1 RNA (**B**). Student's *t-test* was used to compare the differences. *p=0.0277 for comparison of the levels of Dicer1 mRNA in CNs, *p=0.0387 for comparison of Dicer1 RNA in HNs in A**.** *p=0.0469 for comparison of the levels of B1 RNA in CNs, *p=0.0252 for comparison of B1 RNA in HNs**.** The results were averaged from three independent culture preparation.

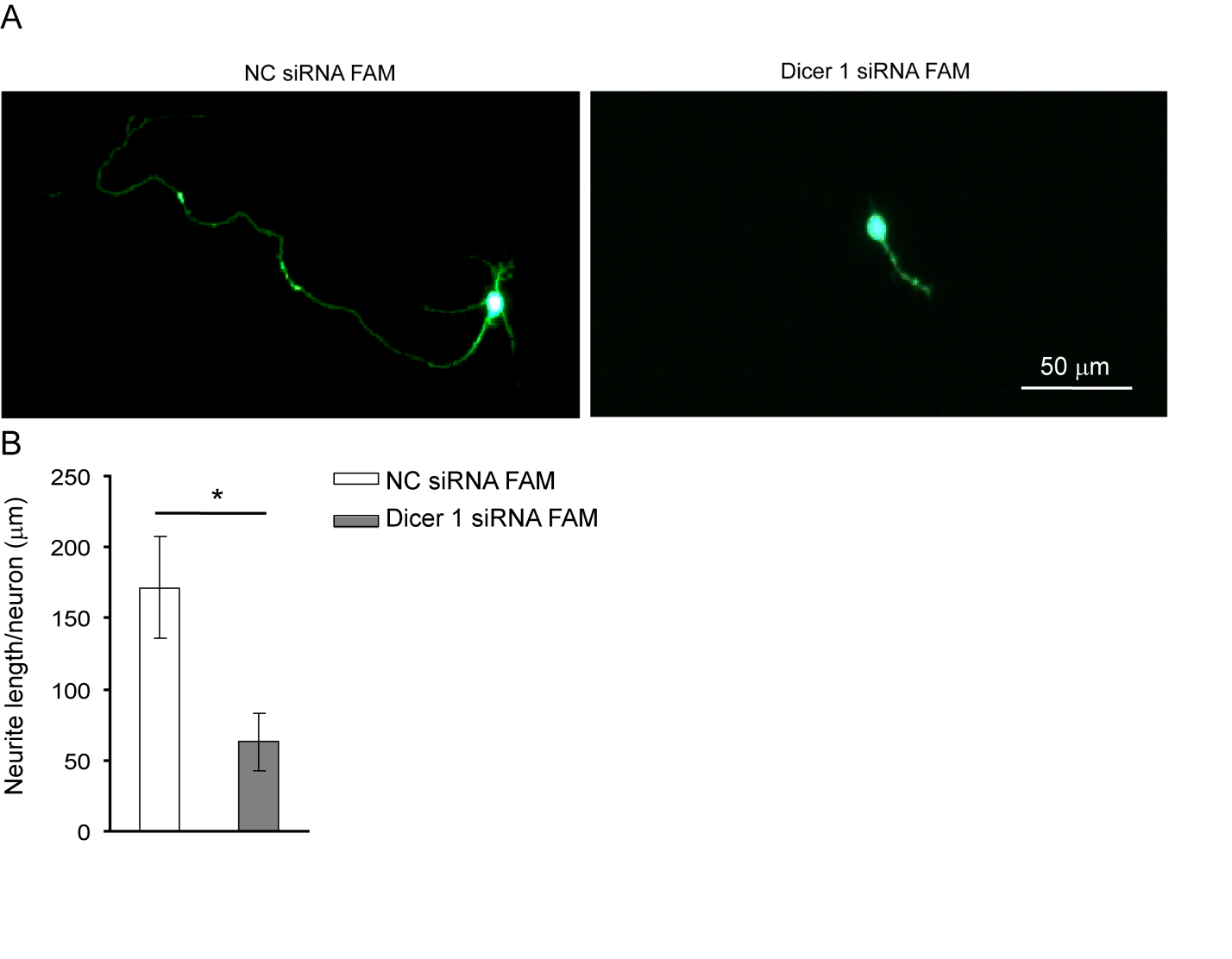

**Figure S3. Knockdown of Dicer1 reduced neurite length in primary cortical neuronal cultures.** 7×10^4^ cortical neurons were transfected with 6-fluorescein amidite (FAM)-labeling Dicer1 siRNA duplex (Dicer1 siRNA FAM) or scrambled siRNA duplex (NC siRNA FAM) for 12 h, respectively. Neurons were observed with fluorescence microscope. **(A)** Representative images of neurons transfected with Dicer1 siRNA FAM or NC siRNA FAM. Bar, 50 μm. **(B)** Total neurite length per cortical neuron averaged from 17 neurons under either transfection condition. Student's *t-test* was used to compare the difference, * p=0.011.

**Supplemental materials and methods**

**Quantitative real-time polymerase chain reaction (qRT-PCR)**

Primary cultured cortical neurons and hippocampal neurons were plated on 6-well plate and treated with Aβ42 oligomer (100 nM) or subject to sham treatment for 48 h in neurobasal medium with 2% B27, respectively. Total RNA were extract by a RNA extracted kit (TIGEN BIOTECH, Co, LTD., Beijing, China, cat#DP424). The following primers were used in qRT-PCR: *Dicer 1* sense, 5'-GTCAGCCGTCAGAACTCACTC-3'; anti-sense, 5'-ACAGTCAAGGCGACATAGCAA-3'. *B1* RNA, sense, 5'-TGCCTTTAATCCCAGCACTT-3'; anti-sense, 5'- GCTGCTCACACAAGGTTGAA-3'. The 18srRNA: sense, 5'-TTCGTATTGCGCCGCTAGA-3', antisense, 5'- CTTTCGCTCTGGTCCGTCTT-3'. The SYBR^TM^ Green (Invitrogen) were used as the dye for qRT-PCR reaction following the manufacturer's instruction. 2^-△△Ct^ method was used to quantify the reaction, which was normalized to sham treatment.
